# Cortical cell ensemble control of past experience-dependent memory updating

**DOI:** 10.1101/2021.07.13.452275

**Authors:** Akinobu Suzuki, Sakurako Kosugi, Emi Murayama, Eri Sasakawa, Noriaki Ohkawa, Ayumu Konno, Hirokazu Hirai, Kaoru Inokuchi

## Abstract

When processing current sensory inputs, animals refer to related past experiences. Current information is then incorporated into the related neural network to update previously stored memories. However, the neuronal mechanism underlying the impact of memories of prior experiences on current learning is not well understood. Here, we found that a cellular ensemble in the posterior parietal cortex (PPC) that is activated during past experience mediates an interaction between past and current information to update memory through a PPC-anterior cingulate cortex circuit in mice. Moreover, optogenetic silencing of the PPC ensemble immediately after retrieval dissociated the interaction without affecting individual memories stored in the hippocampus and amygdala. Thus, a specific subpopulation of PPC cells represents past information and instructs downstream brain regions to update previous memories.

## Introduction

Animals compare current experiences, either implicitly or explicitly, with prior experiences (Dew and Cabeza, 2011). New information is then incorporated into the related neural network to update previously stored memories. Memories are stored in neuronal subpopulations called engrams or memory traces that are activated during experience (Josselyn and Tonegawa, 2020). Memory is then updated through a dynamic process depending on the new situation. For example, original memories can be updated with new information (integration and association) (Schlichting and Preston, 2015) or altered by opposite information (extinction and counterconditioning) (Lee et al., 2017) for survival and higher-order function. Recent studies have begun to clarify the mechanisms underlying memory engrams and how prior experiences affect later learning. Memories encoded close in time interact to generate an associative memory or facilitate the encoding of the later event (Josselyn and Tonegawa, 2020; Nomoto and Inokuchi, 2018). Sharing of engram cell populations mediates the interaction between the prior and the later memories (Abdou et al., 2018; Cai et al., 2016; Nomoto et al., 2016; Rashid et al., 2016; Yokose et al., 2017). These studies have focused on engrams in limbic system structures such as the hippocampus and amygdala, which are primary storage sites of engrams. Human studies have indicated that a cortical modulation system controls the active integration of new information into the neural network that represents past information (Hasson et al., 2015; Parsons, 2018; Schlichting and Preston, 2015). Thus, cortical top-down modulation may instruct engrams stored in the downstream regions to interact to update memory.

The posterior parietal cortex (PPC) is a major associative region in the mammalian cortex. The PPC is classically considered to be associated with cognitive functions, such as visual perception (Pisella et al., 2009), spatial attention and visual guidance (Driver et al., 2010; Husain and Nachev, 2007), and episodic memory in human and rodent (Berryhill et al., 2007) (Jonker et al., 2018) (Myskiw and Izquierdo, 2012). Recent findings indicate that the PPC plays important roles in memory updating. The PPC modulates fear memory renewal after the extinction in fear conditioning (Joo et al., 2020). The PPC represents past sensory history and influences behavioral outcome upon current experience (Akrami et al., 2018). Furthermore, the PPC implements updating of the prediction when the uncertainty of the prediction decreases with new sensory inputs (Funamizu et al., 2016). However, an important and outstanding question is how representations of past experiences in the PPC interact with current sensory inputs. The complexity of the experimental design used in the previous studies makes it technically difficult to identify the specific cellular populations and assign them to past or current experience. In this study, by taking advantage of a simple behavioral task in which mice associate a past experience (context) with a current sensory input (footshock) during one learning session, we clarified the interaction mechanism at the cell population level.

## Results

### Neuronal ensemble in PPC responds to past and current experiences

In a context pre-exposure and immediate shock (IS) task, animals retrieve a previously encoded memory of the pre-exposed context, which is then associated with the current experience, an IS in the same context, to update memory, such that animals exhibit a fear response in this context (Fanselow, 1990; Ohkawa et al., 2015). This is followed by an increase in the degree of overlapping in engram cell populations both in the hippocampus and amygdala (Ohkawa et al., 2015). This is a simple learning task, and so is suitable for detecting the corresponding neuronal ensembles responsible for memory updating.

We found that an IS without pre-exposure failed to associate the context and shock (Figures 1A and 1B). Conversely, mice showed higher freezing in test sessions when pre-exposed and IS contexts were the same (paired) than distinct (unpaired) contexts, irrespective of the interval between context pre-exposure and the IS (Figures 1B and 1C). Cellular compartment analysis of temporal activity by fluorescence in situ hybridization (CatFISH) utilizing an immediate early gene Arc permits the detection of activity-dependent activation of large neuronal ensembles during two distinct events(Guzowski et al., 1999). After neuronal activation, Arc RNA is initially localized to the nucleus but then translocates to the cytoplasm within 30 min. Therefore, the expression of cytoplasmic and nuclear Arc RNA can be used to distinguish neuronal populations engaged by the behavioral experience of the first event (pre-exposed context) from those engaged by the behavioral experience of the second event (IS). CatFISH revealed that, in addition to the hippocampus and amygdala, several cortical regions responded to both events (Figure S1). The neuronal population of cytoplasmic and nuclear Arc double-positive cells in the posterior parietal cortex (PPC) of the paired group was significantly larger than that in the unpaired and No IS groups (Figures 1D-1F). This suggests that the neuronal population of double-positive cells in the PPC responds to past and current experiences. The degree of overlap in each group was significantly higher than the corresponding chance level. Moreover, the degree of Arc double-positive cells of the No IS group was significantly larger than that in the unpaired group. This suggests that memory updating might take place even when the behavioral phenotype does not change significantly.

**Figure 1.**
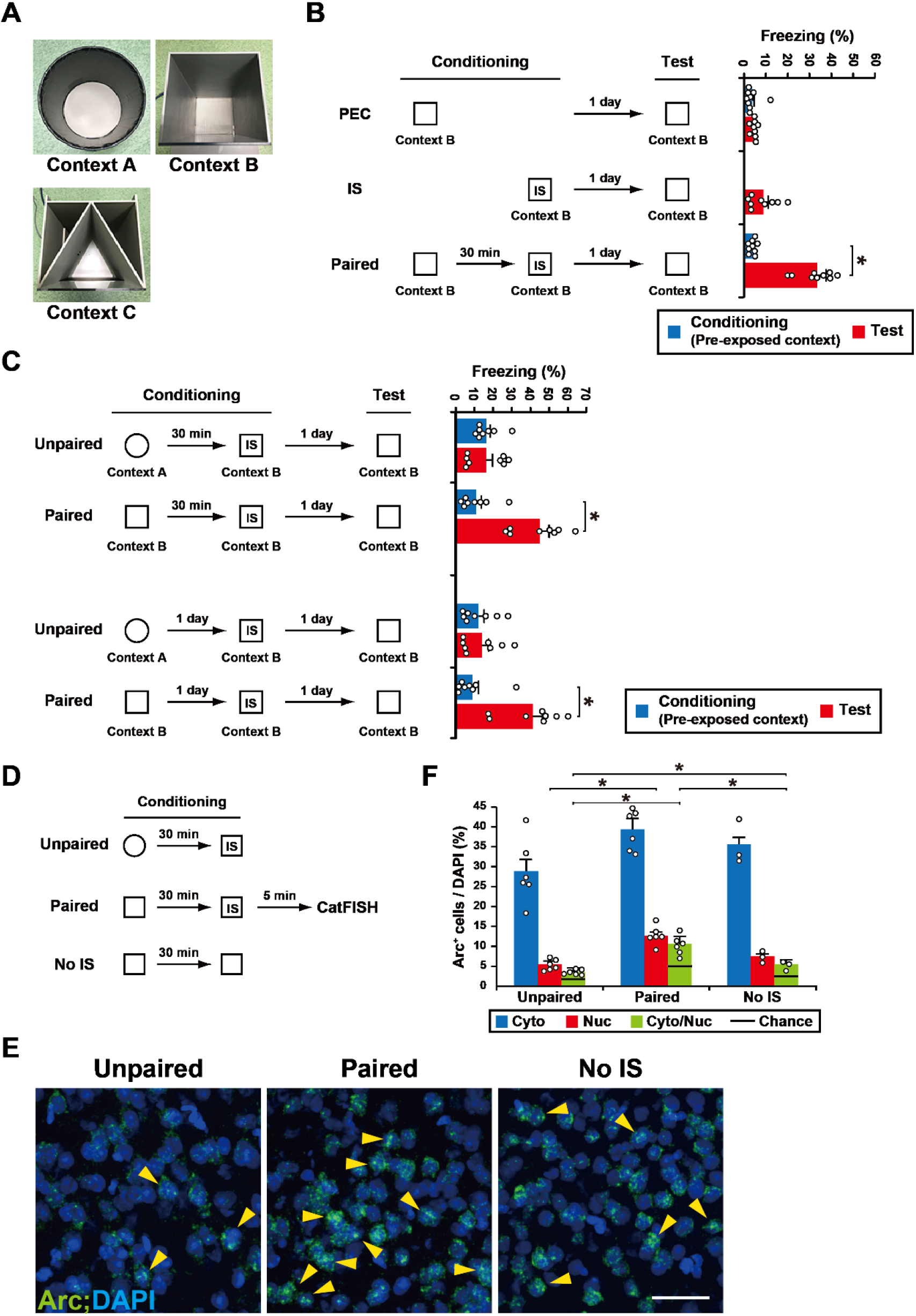
PPC cell ensembles corresponding to the pre-exposed context and IS. (A) Photo showing context A, B, and C used in this study. (B) Schema of pre-exposed context (PEC), immediate shock (IS), and PEC-IS paired paradigm. The graph shows the freezing levels during the conditioning and test session (n = 9 mice/group). (C) Schema of the unpaired and paired paradigm. Mice were pre-exposed to context A (unpaired group) or context B (paired group), and then received an IS in context B 30 min or 1 day later. Mice were then tested 1 day after the conditioning. The graph shows the freezing level observed during the conditioning and test session (interval between Ctx and IS: 30 min: n = 8 mice/group; 1 day: n = 8 mice/group). (D) Schema of Arc CatFISH analysis. Three groups (unpaired, paired, and the No IS group, which did not receive a footshock) were used in this study. Mice were sacrificed for Arc CatFISH analysis 5 min after the conditioning. (E) Representative images of the PPC CatFISH analysis. The Arc RNA signal and DAPI nuclear staining are shown in green and blue, respectively. The yellow arrowheads indicate Nuclear+, Cytoplasmic+ (double positive) cells. Scale bar, 50 μm. (F) Percentage of Arc+ cells per DAPI+ cells (unpaired and paired group, n = 6 mice, No IS group, n = 3 mice). Error bars indicate the mean ± s.e.m. *P < 0.05. Cyto, cytoplasmic; Nuc, nuclear. Black line indicates chance. For results of other brain regions and details of statistical data, see Figure S1 and Table S1.

### Optical silencing of neuronal ensemble in PPC blocks memory association

We used c-fos::tTA mice that express the tetracycline-transactivator (tTA), which is under the control of c-fos promoter, an immediate early gene used as a marker of neuronal activity (Figure 2A). In the absence of doxycycline (Dox), activation of the c-fos promoter leads to tTA expression, and tTA binding to the tetracycline-responsive element (TRE) induces target gene expression. In the presence of Dox, tTA is prevented from binding to the TRE. We found that context pre-exposure induced tTA mRNA expression in the PPC (Figure S2A). A recombinant lentivirus (LV) encoding TRE::ArchT-EYFP or TRE::EYFP was injected bilaterally into the PPC of c-fos::tTA mice to label cells that were activated during context pre-exposure (Figures 2B and S2B). LV-injected mice were taken off Dox (OFF Dox group) for 2 days, and then exposed to the context B to label ArchT-EYFP or EYFP proteins in neurons that were activated during pre-exposure. The next day, mice received an IS in the same context while optical illumination (continuous 589 nm light) was delivered to the PPC during the IS session (Light ON). The function of ArchT-EYFP was restricted to the Light ON/OFF Dox group; the IS session-induced expression of Zif268 (also called Egr1), an immediate early gene, in the PPC was specifically suppressed in this group (Figure S3). The other groups were treated in the same way. The ArchT-Light ON group showed significantly lower freezing in the test session than the ArchT-Light OFF and EYFP-Light ON groups (Figure 2B). These results indicate that a PPC neuronal population is required for the association between the pre-exposed context and IS. Furthermore, the silencing of PPC cells that were activated in a different context had no effect on fear expression in the pre-exposed context. PPC cells, which were activated during pre-exposure to the circle context, were labeled with ArchT (circle-labeled group). Mice received the paired condition of the CPFE paradigm with optical silencing to the PPC during an IS session in the square context. The circle-labeled group showed significantly higher freezing in the test session even when optical silencing was delivered to the PPC cell population that responded to the circle context (Figure 2C). This indicates that cell populations labeled in the PPC are specific to each context.

**Figure 2.**
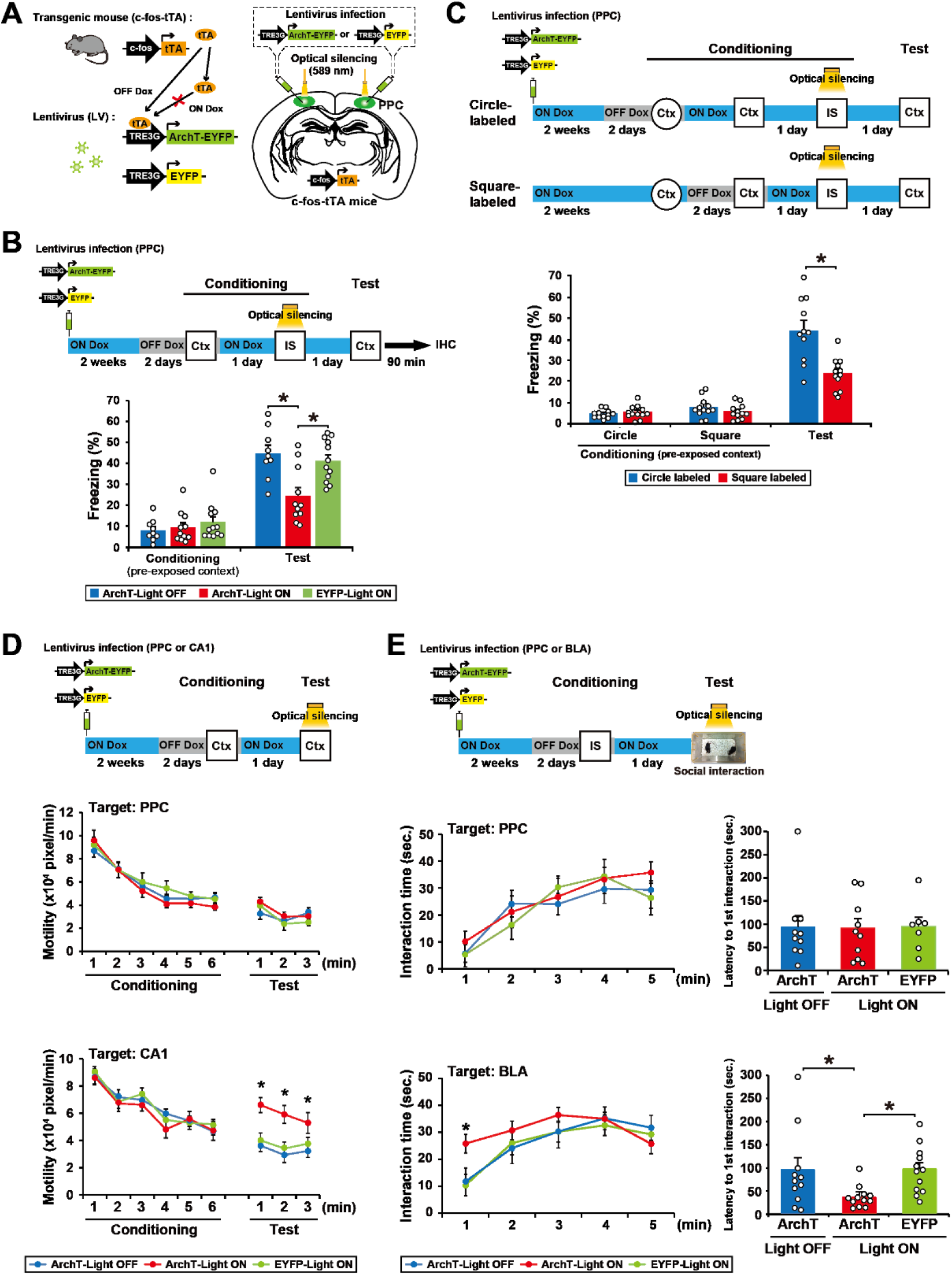
Optical silencing of cell assemblies in the PPC blocks memory association without affecting the original memories. (A) Labeling of PPC neurons in c-fos::tTA transgenic mice with the LVs TRE3G::ArchT-EYFP or TRE3G::EYFP. (B) Silencing of the cell ensembles during an IS session (top). The blue and gray bars indicate the presence or absence of Dox, respectively. The graph shows the freezing level of each group during the conditioning and test session (bottom) (ArchT-Light OFF, n = 9 mice, ArchT-Light ON, n = 11 mice, EYFP-Light ON, n = 12 mice). Mice were sacrificed 90 min after the test for immunohistochemistry (Figures 6A and 6B). (C) Silencing of the cell ensembles corresponding to circle or square context during an IS session (top). The graph shows the freezing level of each group during the conditioning and test session (bottom) (Circle-labeled, n = 12 mice, Square-labeled, n = 12 mice). (D) Optical silencing of context exposure-activated cell ensembles in PPC or CA1 during the test session (top). The graph shows motility during the conditioning and test sessions in the groups with silencing of cell ensemble in PPC (middle) and CA1 (bottom) (PPC; ArchT-Light OFF, n = 10 mice, ArchT-Light ON, n = 10 mice, EYFP-Light ON, n =7 mice, CA1; ArchT-Light OFF, n = 12 mice, ArchT-Light ON, n = 12 mice, EYFP-Light ON, n = 13 mice). (E) Silencing the IS session-activated cell ensembles in PPC or BLA during the social interaction test (top). The graph shows the interaction time (left) and latency to the first interaction (right) during the test session in the groups with silencing cell ensemble in PPC (middle) and BLA (bottom) (PPC; ArchT-Light OFF, n = 10 mice, ArchT-Light ON, n = 10 mice, EYFP-Light ON, n = 7 mice, BLA; ArchT-Light OFF, n = 11 mice, ArchT-Light ON, n = 12 mice, EYFP-Light ON, n = 12 mice). Error bars indicate the mean ± s.e.m. *P < 0.05. Ctx, context; IS, immediate shock; IHC, immunohistochemistry. See also Figure.S2, S3 and S4. For details of statistical data, see Table S1.

Optical silencing of the PPC cells activated during the context pre-exposure had no effect on the retrieval of pre-exposed context memory (the ArchT-Light ON group), which was evident from the finding that motility, an indicator of contextual memory (Kitamura et al., 2012), was comparable to other groups (Figure 2D). This result suggests that PPC is not required when there is no memory updating. When the same strategy was applied to the hippocampal CA1 region, the ArchT-Light ON group showed significantly higher motility than the other groups, which indicates that CA1 acts as a memory storage site of the pre-exposed context (Figure 2D).

We applied the social interaction test to measure the IS memory recall under the assumption that IS-induced fear would suppress social interaction behavior. Indeed, mice that received IS showed a decrease in interaction time and an increase in latency to first interaction compared with No IS mice (Figure S2C). Thus, the social interaction test is appropriate for the assessment of whether or not IS memory was intact. Optical silencing of basolateral amygdala (BLA) neurons that were activated during the IS enhanced social interaction behavior, with a longer interaction time and shorter latency compared with the control groups (Figure 2E). By contrast, optical silencing of the PPC cells activated by the IS did not affect social interaction, which was not significantly different from the other groups (Figure 2E). This finding indicates that IS memory was stored in the BLA and IS memory recall was independent from the activity of the PPC cell ensemble. Similar results were obtained when the sodium channel blocker lidocaine was injected into the PPC (Figure S4). This treatment inhibited the pre-exposed context-IS association (Figure S4A) without affecting the proper recall of the pre-exposed context and IS memories (Figures S4B and S4C; compare to vehicle control).

### Optical activation of neuronal ensemble in PPC generates an artificial associative memory

We next examined whether stimulation of the PPC cell ensemble can generate an artificial pre-exposed context-IS associative memory (Figure 3). We applied an unpaired paradigm in which mice did not associate the pre-exposed context with the IS. ChR2-EYFP functioned in the PPC cell ensemble specifically in the Light ON and OFF Dox condition, which was judged by endogenous c-Fos induction (Figures 3A and S5). The c-fos::tTA mice who received LV vector injections into PPC were in the OFF Dox condition for 2 days, and then pre-exposed to context A to label the activated PPC cells. One day later, mice received an IS in the different context B while 20 Hz light pulses were delivered to the PPC. The ChR2-Light ON group exhibited a higher freezing response in the pre-exposed context (context A), in which mice did not receive a footshock, than the other groups (Figure 3B). The freezing response of the ChR2-Light ON group in the neutral context (context C) was comparable to that of the control group, which indicates that optically-induced artificial memory is context-specific.

**Figure 3.**
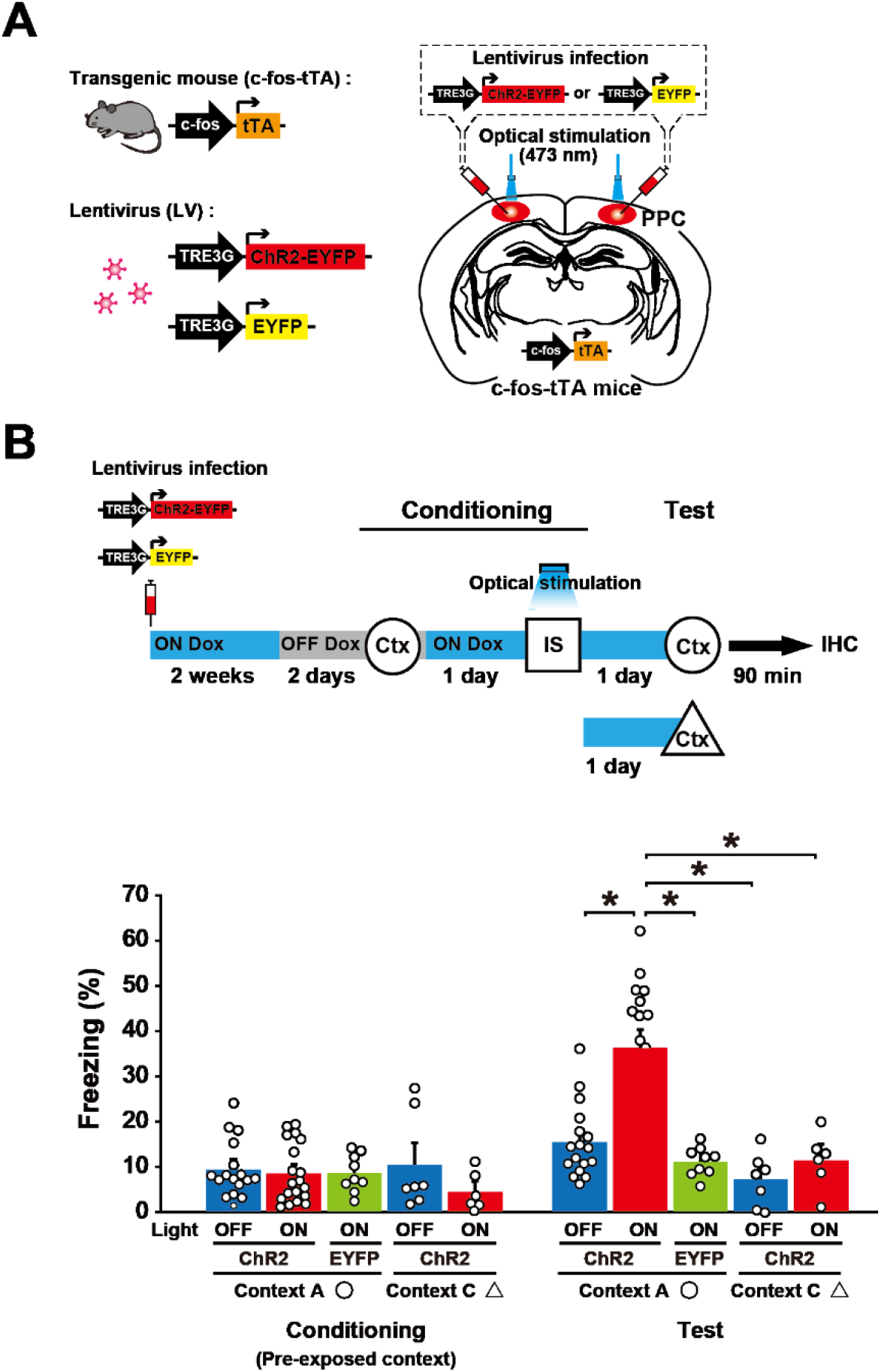
Optical activation of the PPC cell ensemble generates an artificial Pre-exposed context-IS associative memory. (A) Labeling of the PPC neurons in c-fos::tTA transgenic mice with the LVs TRE3G::ChR2-EYFP or TRE3G::EYFP. (B) Activating cell ensembles during an IS session of conditioning (top). The blue and gray bars indicate the presence or absence of Dox, respectively. The graph shows freezing level in context A or C in the conditioning and test session (bottom) (ChR2-Light OFF in circle, n = 17 mice, ChR2-Light ON in circle, n = 19 mice, EYFP-Light ON in circle, n = 9 mice, ChR2-Light OFF in triangle, n = 7 mice, ChR2-Light ON in triangle, n = 6 mice). Mice were sacrificed 90 min after the test for immunohistochemistry (Figures 6C and 6D). Error bars represent the mean ± s.e.m. *P < 0.05. Ctx, context; IS, immediate shock; IHC, immunohistochemistry. See also Figure S5. For details of statistical data, see Table S1.

### The PPC regulates associative memory retrieval

Suppression of the PPC ensemble activity transiently dissociated the pre-exposed context-IS association that had already been stored in the brain (Figures 4A and 4B). A subset of the PPC cell ensemble that responded to memory reactivation was labeled with ArchT-EYFP. Optical silencing of the PPC ensemble during test 1 suppressed memory recall. However, the suppressed memory returned to the control level the next day (test 2).

**Figure 4.**
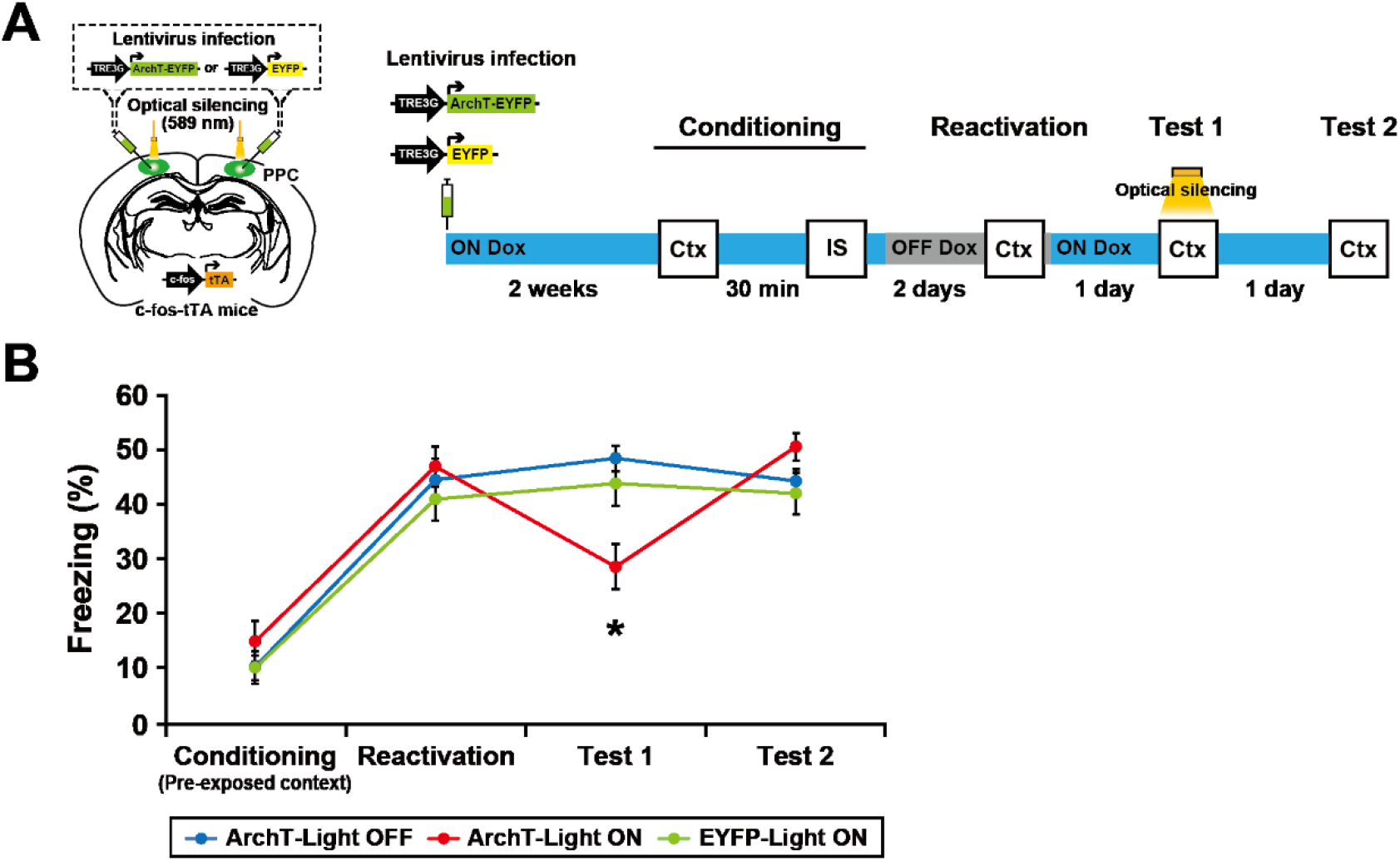
The PPC regulates associative memory retrieval. (A) Schematics showing labeling of the PPC neurons in c-fos::tTA transgenic mice with the LVs TRE3G::ArchT-EYFP or TRE3G::EYFP (left). Schematic of the behavioral experiment with optical silencing (right). Blue and gray bars indicate the presence or absence of Dox, respectively. Optical silencing to the PPC was delivered during test 1. (B) The graph shows the freezing level during the conditioning, reactivation, test 1, and test 2 (ArchT-Light OFF, n = 14 mice, ArchT-Light ON, n = 13 mice, EYFP-Light ON, n = 10 mice). Error bars represent the mean ± s.e.m. *P < 0.05. Ctx, context; IS, immediate shock. For details of statistical data, see Table S1.

### Optical silencing of PPC ensemble activity dissociates the memory association

A previous study indicated that optical inhibition of CA1 neurons for 15 min immediately after memory reactivation leads to disruption of fear memory (Lux et al., 2017). Consistent with previous evidence, optical silencing of PPC neurons immediately after test 1 suppressed the fear memory recall 1 day later in test 2 (Figures 5A and 5B), presumably via a reconsolidation inhibition-like mechanism. In the other groups, optical silencing had no effect on recall of pre-exposed context (Figure 5C, motility, test 2) or IS (Figure 5D, vehicle, test 3) memories. When CA1 activity was suppressed by lidocaine injection, mice showed a higher motility in test 3 than vehicle-injected mice (Figure 5C). Similarly, BLA suppression resulted in a longer interaction time and shorter latency of the first interaction (Figure 5D, test 3, lidocaine). Thus, suppression of pre-exposed context-IS associative memory (Figure 5B) was not due to an impairment in pre-exposed context or IS memory.

**Figure 5.**
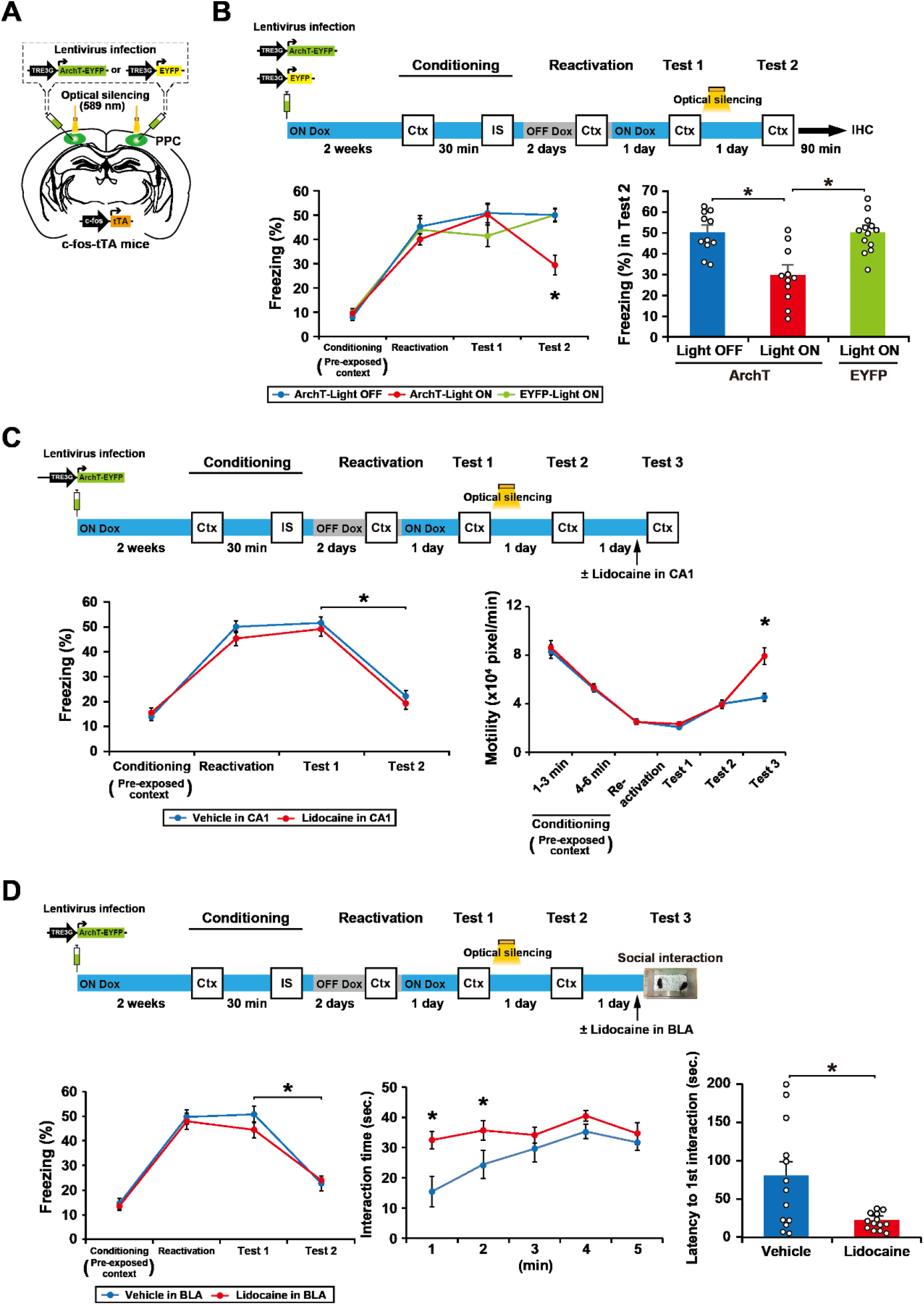
Optical silencing of PPC ensemble activity dissociates the pre-exposed context-IS association without affecting original memories. (A) Labeling of PPC neurons in c-fos::tTA transgenic mice with the LVs TRE3G::ArchT-EYFP or TRE3G::EYFP. (B) The behavioral experiment with optical silencing (top). Blue and gray bars indicate the presence or absence of Dox, respectively. Optical silencing (15 mins) to the PPC was delivered in the home cage immediately after test 1. The left graph shows the freezing level at conditioning, reactivation, test 1, and test 2 (bottom) (ArchT-Light OFF, n = 11 mice, ArchT-Light ON, n = 11 mice, EYFP-Light ON, n = 13 mice). The right graph shows freezing level of each group at test 2. Mice were sacrificed 90 min after test 2 for immunohistochemistry (Figures 6E and 6F). (C) The behavioral experiment with 15 min optical silencing (top). Optical silencing to the PPC was delivered in the home cage immediately after test 1. Lidocaine was injected in the CA1 region 15 mins before test 3. The graph shows the freezing level at conditioning, reactivation, test 1, and test 2 (left), and motility during the conditioning, reactivation, test 1, test 2 and test 3 (right) (Vehicle in CA1, n = 16 mice, Lidocaine in CA1, n = 17 mice). (D) The behavioral experiment with 15 min optical silencing (top). Optical silencing to the PPC was delivered in the home cage immediately after test 1. Lidocaine was injected in the BLA 15 min before test 3. The graph shows the freezing level at conditioning, reactivation, test 1, and test 2 (left), the interaction time (center) and the latency to the first interaction (right) in test 3 (bottom) (Vehicle in BLA, n = 13 mice, Lidocaine in BLA, n = 13 mice). Error bars represent the mean ± s.e.m. *P < 0.05. Ctx, context; IS, immediate shock; IHC, immunohistochemistry. For details of statistical data, see Table S1.

### Optical silencing or activation affects c-Fos expression in the BLA and ACC after a test session

What neural circuit downstream of the PPC regulates pre-exposed context-IS memory association? We found a decrease or increase, respectively, in the number of c-Fos-positive cells when the PPC cells of interest were optically silenced (Figures 6A, 6B, 6E and 6F; paired condition) or stimulated (Figures 6C and 6D; unpaired condition), not only in the BLA, but also in the anterior cingulate cortex (ACC), which also plays a critical role in regulating fear responses (Allsop et al., 2018; Etkin et al., 2011; Gross and Canteras, 2012).

**Figure 6.**
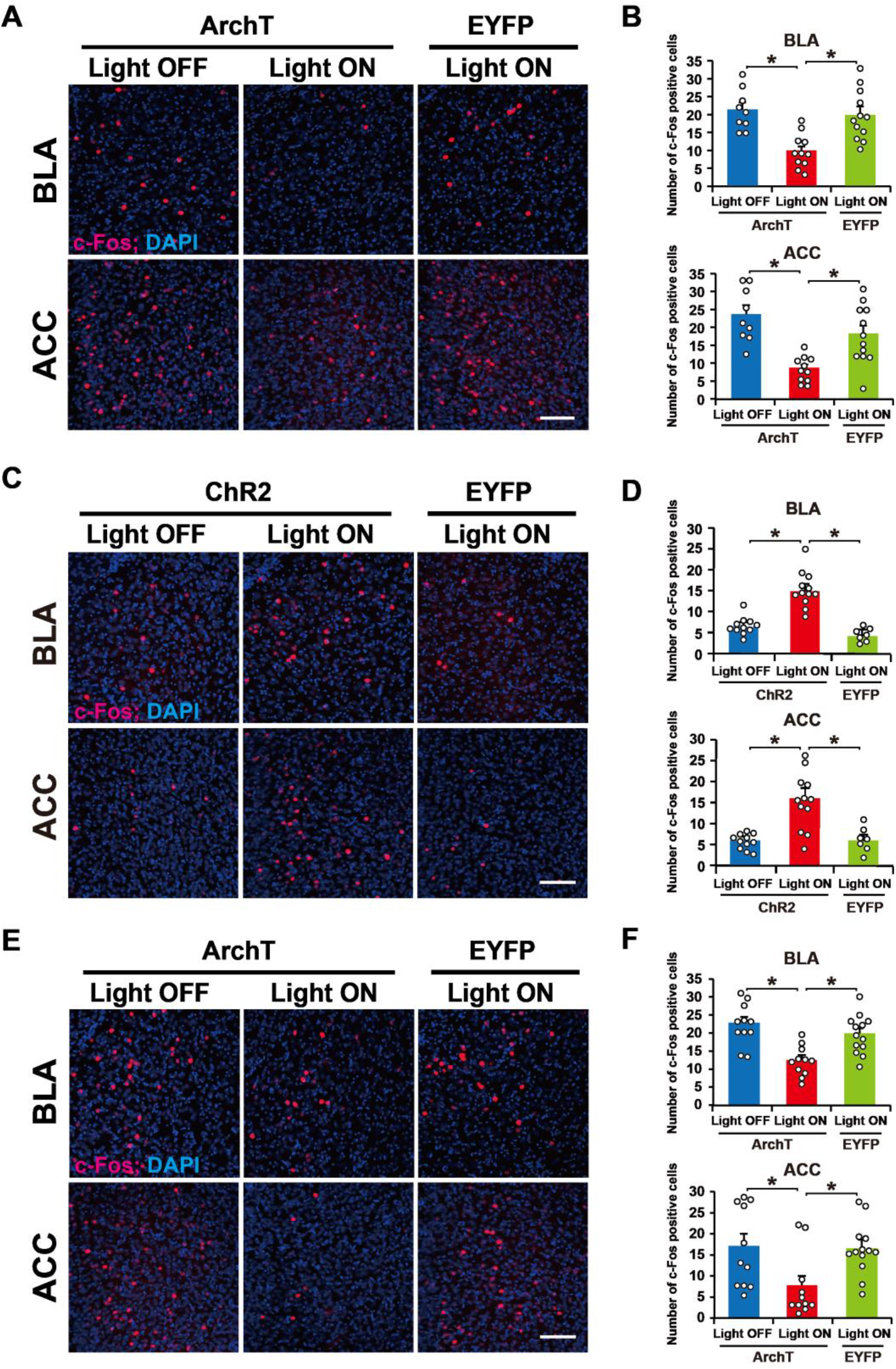
c-Fos-positive cells in the BLA and the ACC after the test session. (A, C and E) Representative images of c-Fos expression 90 min after the test shown in Figure 2B (panel A), Figure 3B (panel C), and Figure 5B (panel E). Scale bar, 100 μm. (B, D and F) Quantitative proportion of c-Fos-positive cells in the BLA (top) and ACC (bottom) (panel B: ArchT-Light OFF, n = 9 mice, ArchT-Light ON, n = 11 mice, EYFP-Light ON, n = 12 mice; panel D: ChR2-Light OFF, n = 11 mice, ChR2-Light ON, n = 12 mice, EYFP-Light ON, n = 7 mice; panel F: ArchT-Light OFF, n = 11 mice, ArchT-Light ON, n = 11 mice, EYFP-Light ON, n = 13 mice.) Error bars indicate the mean ± s.e.m. *P < 0.05. For details of statistical data, see Table S1.

### PPC-ACC circuit regulates memory association

The PPC was found to be bidirectionally connected to the ACC by injecting an anterograde tracer, Biotinylated Dextran Amines (BDA), into the PPC and a retrograde tracer, cholera toxin subunit B-Alexa Fluor 488 conjugate (CTB488), into the ACC (Figures 7A, S6A and S6B). To examine the involvement of this PPC-ACC circuit in memory association, we performed a similar experiment to that shown in Figure 2B, except that AAV9 TRE::ArchT-EYFP was infused directly into the PPC and optical silencing was delivered to the ACC (Figures 7B and S6C). The expression and function of ArchT-EYFP in the ACC was restricted to the Light ON and OFF Dox condition (Figure S7). Optical silencing of the PPC neuronal terminals at the ACC that was delivered during the IS session blocked the formation of memory association (Figure 7C). This was accompanied by a decrease in the number of c-Fos-positive cells in the BLA and ACC (Figure 7D), indicating that the PPC-ACC circuit regulates memory updating.

**Figure 7.**
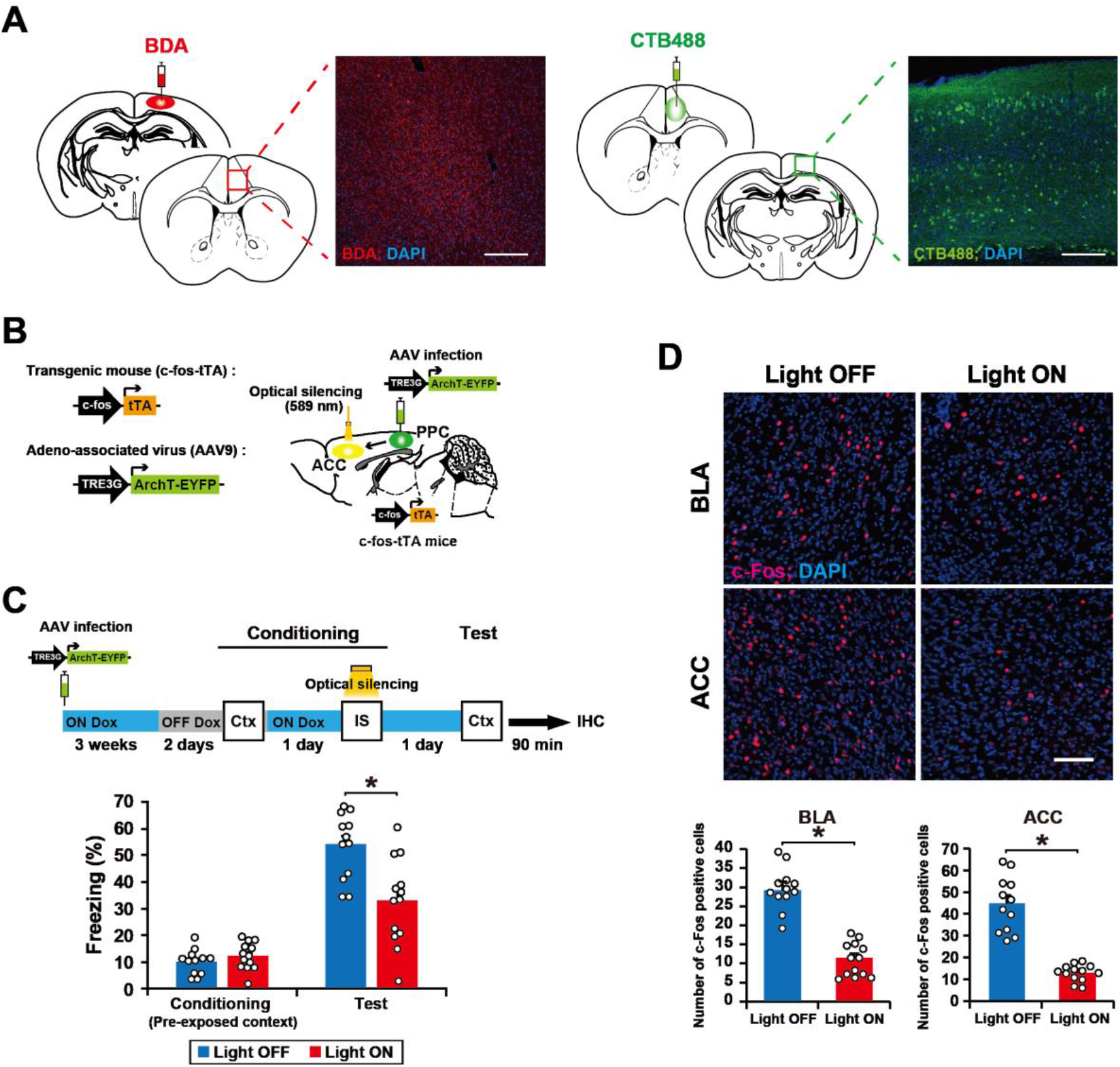
A PPC-ACC circuit regulates memory association. (A) Horizontal brain sections of the ACC from a mouse injected with BDA in the PPC, in which neurons were immunostained with streptavidin-Alexa Fluor 555 (red; left). PPC cell axons were detected in the ACC. Horizontal brain sections of the PPC from a mouse injected with CTB488 in the ACC (green; right). CTB488-labeled cells were detected in the PPC. Scale bar, 200 μm. (B) Schematics showing labeling and manipulation of the PPC-ACC circuit in c-fos::tTA transgenic mice with the AAV9 TRE3G::ArchT-EYFP. (C) Silencing of cell ensembles during an IS session of conditioning (top). Blue and gray bars indicate the presence or absence of Dox, respectively. Mice were sacrificed 90 min after the test for immunohistochemistry (panel D). The graph shows the freezing level at the conditioning and test session (bottom) (Light OFF, n = 12 mice, Light ON, n = 13 mice). (D) Representative image of c-Fos expression 90 min after the test (top). Scale bar, 100 μm. The proportion of c-Fos-positive cells in the BLA (left) and ACC (right) (Light OFF, n = 12 mice, Light ON, n = 13 mice). Error bars represent the mean ± s.e.m. *P < 0.05. Ctx, context; IS, immediate shock; IHC, immunohistochemistry. See also Figure S6 and S7. For details of statistical data, see Table S1.

## Discussion

Our study reveals that a PPC cell ensemble recruited during a past experience acts as a top-down modulator for the interaction between recalled memory and ongoing experience. This is consistent with previous findings that the PPC represents past experiences (Akrami et al., 2018) and is involved in episodic memory retrieval (Sestieri et al., 2017). The PPC has been proposed to act as sensory history buffer, i.e., past experience is held in memory for later comparison with present experience (Bitzidou et al., 2018). Our findings suggest that the PPC cell ensemble that is activated during past experience (context pre-exposure) acts as a sensory buffer for use in a future relevant experience.

The PPC ensemble identified shares characteristics with the memory engram (Josselyn and Tonegawa, 2020), in that activity in both is generated as a cellular ensemble during experience. Furthermore, memories can be modified by manipulating the activity of the PPC ensemble, which is similar with previous studies where manipulation of engram cells in the hippocampal and amygdala generates false memories (Ohkawa et al., 2015; Ramirez et al., 2013). However, our results indicate that PPC ensemble function is distinct from that of the memory engram because silencing PPC ensemble activity had no effect on the recall of individual memories of context and fear. Thus, the cell ensemble in the PPC does not store pre-exposed context and IS memories but rather regulates the interaction between these memories, i.e., when original memory is updating. Our previous study showed that the hippocampus and BLA ensembles serve as an engram for pre-exposed context and IS, respectively (Ohkawa et al., 2015). Moreover, neocortical memory engram is thought to form slowly in systems memory consolidation (Frankland and Bontempi, 2005). Indeed, it has been reported that memory engram cells in the neocortex are generated during learning in a silent form, and then gradually mature to form a functional engram (Kitamura et al., 2017; Tonegawa et al., 2018). By contrast, we found that PPC ensemble cells quickly become functional after learning, which is consistent with recent findings of a rapid involvement of neocortical areas in memory processing (Brodt et al., 2018; Hebscher et al., 2019). Our findings suggest that these PPC cells may recruit engram cells in the downstream regions, i.e., the hippocampus and amygdala, upon presentation of the context to express fear memory. The immunohistochemical results (Figure 6) support this notion by showing that manipulation of the PPC ensemble regulates the number of c-Fos-positive cells in the BLA.

We found that the PPC ensemble is required not only for encoding, but also for the recall of associated memory. Post-retrieval suppression of the corresponding cell ensemble dissociated the linkage between pre-exposed context and IS without affecting individual memories. Taken together, our findings indicate that the PPC ensemble can flexibly bridge and disconnect different kinds of information, namely the pre-exposed context and IS, that is stored in downstream regions in a top-down manner.

Another important issue is the identity of the PPC-connected brain region that mediates the role of the PPC in memory updating. We found that the PPC has reciprocal connections with the ACC and that optical silencing of the axon terminal of the PPC cell ensemble in ACC during the IS session blocked the association between pre-exposed context and IS memories. Moreover, the PPC ensemble recruited ACC and BLA neurons (Figure 6). ACC neurons project to the BLA (Gross and Canteras, 2012), and the PPC has reciprocal connections with the hippocampus via entorhinal and retrosplenial cortexes (Save and Poucet, 2009). The medial prefrontal cortex, including the ACC, is involved in encoding novel but related information into existing knowledge in human and existing schema in rodent (Gilboa and Marlatte, 2017; Sommer, 2016; Tse et al., 2007; Tse et al., 2011). Although further study is required, the PPC ensemble may, via the ACC and entorhinal and retrosplenial cortexes, instruct downstream regions (hippocampus and amygdala) to integrate newly acquired information into a related neural network that represents past experiences to create sematic memories and schema.

Re-experiencing unwanted traumatic events, i.e., flashbacks, is a major symptom of post-traumatic stress disorder that is triggered by stimuli that were present around the traumatic experience, most of which relate in some way to daily events (Ehlers, 2010; Lissek and van Meurs, 2015). Thus, our findings may inform research that aims to prevent intrusion-based flashbacks in patients with post-traumatic stress disorder.

## Acknowledgments

We thank K. Deisseroth (Stanford University) for the ArchT 3.0-EYFP cDNA and ChR2(T159C)-EYFP cDNA; H. Hioki and T. Kaneko (Kyoto University) for the TGB vector; H. Nomura (Hokkaido University) for guidance with CatFISH method; H. Nishizono, M. Matsuo, and Y. Arasaki-Yanagihashi for generating the transgenic mice; and S. Tsujimura and S. Okami for maintaining the transgenic mice. All members of the Inokuchi laboratory supported and discussed this study. This work was supported by the Core Research for Evolutional Science and Technology (CREST) program (JPMJCR13W1) of the Japan Science and Technology Agency (JST), JSPS KAKENHI grant numbers JP18H05213 and JP23220009, a Grant-in-Aid for Scientific Research on Innovative Areas “Memory dynamism” (JP25115002) from MEXT, and the Takeda Science Foundation to K.I.; JSPS KAKENHI JP 24680034, the Hokuriku Bank Grant for Young Scientists, The Uehara Memorial Foundation, The Takeda Science Foundation, The Kanae Foundation for the promotion of medical science, and the Tamura Science and Technology Foundation to A.S.; and Brain Mapping by Integrated Neurotechnologies for Disease Studies (Brain/MINDS) from Japan Agency for Medical Research and development, AMED (JP20dm0207057) to H.H.

## Author Contributions

A.S. and K.I. designed the study. A.S., S.K., and E.M. cloned all vector constructs. S.K., E.M., A.K., and H.H. prepared the viruses. N.O. introduced the optogenetics techniques. A.S., S.K., E.M., and E.S performed all animal surgery, behavioral experiments, histology, and data analyses. A.S. and K.I. wrote the manuscript and contributed to the interpretation. K.I. supervised the entire project. All authors discussed and commented on the manuscript.

## Declaration of interests

The authors declare no competing interests.

## Methods

### Animals

All animal procedures were performed in accordance with the guidelines of the National Institutes of Health (NIH) and were approved by the Animal Care and Use Committee of the University of Toyama. Naïve male C57BL/6J (Japan SLC, Inc., Shizuoka, Japan) and c-fos::tTA transgenic mice (Mutant Mouse Regional Resource Centre, stock number: 031756-MU) were purchased as described previously (Ohkawa et al., 2015). c-fos::tTA transgenic mice were raised from the time they were fetuses on food containing 40 mg/kg Dox and maintained on Dox pellets, except for the labeling day. All mice were maintained on a 12 h light/dark cycle (lights on 8:00 am – 8:00 pm) at 24 ± 3°C and 55 ± 5% humidity, had ad libitum access to food and water, and were housed in a cage with littermates until surgery.

### Viral constructs

The pLenti-TRE3G::eArchT3.0-EYFP and pLenti-TRE3G::EYFP were used as previously described (Nomoto et al., 2016; Yokose et al., 2017). The pLenti-TRE3G::ChR2(T159C)-EYFP plasmid was constructed using a two-step process. The ChR2(T159C)-EYFP fragment of pLenti-CaMKII::ChR2(T159C)-EYFP was prepared with the BamHI-EcoRI restriction site. The EcoRI sites of the ChR2(T159C)-EYFP fragment was blunt ended with the Klenow fragment of Escherichia coli DNA polymerase I (Takara Bio Inc., Shiga, Japan, 2140A). The resulting ChR2(T159C)-EYFP fragment was subcloned into pLenti-TRE3G::IRES at the BglII-EcoRV restriction sites, generating a pLenti-TRE3G::ChR2(T159C)-EYFP plasmid. Then, the TRE3G::ChR2(T159C)-EYFP fragment was prepared with EcoRI-BamHI restriction sites. The BamHI sites of the TRE3G::ChR2(T159C)-EYFP fragment was blunt ended. The resulting TRE3G::ChR2(T159C)-EYFP fragment was subcloned into the EcoRI-EcoRV restriction site of the TGB plasmid (Hioki et al., 2009), generating a pLenti-TRE3G::ChR2-EYFP plasmid. The above lentiviral plasmids were prepared using an EndoFree Plasmid Maxi kit (Qiagen, Hilden, Germany) and were used for the lentivirus preparation as described previously (Ohkawa et al., 2015).

The pAAV-TRE3G::ArchT-EYFP plasmid was constructed using the following steps. The TRE3G fragment of pLenti-CaMKII::ArchT3.0-EYFP was prepared with the XhoI-BamHI restriction site. The XhoI sites of the TRE3G fragment was blunt ended with the Klenow fragment of Escherichia coli DNA polymerase I. The resulting TRE3G fragment was subcloned into the pAAV-CaMKII::ArchT3.0-EYFP plasmid at the MluI-BamHI restriction sites, of which the MluI site was blunt ended, generating a pAAV-TRE3G::ArchT3.0-EYFP plasmid. The recombinant AAV vector was produced using the minimal purification (MP) method(Konno and Hirai, 2020). Six days after the plasmids had been transfected to AAV293 cells, the culture supernatant was collected into a 50 ml tube and then centrifuged at 2,000 rpm for 10 min by centrifuge with swing-rotor (KN-70, level = 7, KUBOTA, Tokyo, Japan). The supernatant was filtered through a Millex-HV 0.45 μm syringe-filter (SLHV033R, Merck Millipore, Darmstadt, Germany) into a new 50 ml tube. The filtered supernatant was poured into the optimized VIVASPIN filter unit (No. VS2041, Sartorius Stedim Lab Ltd., Stonehouse, UK) and was centrifuged at 3,000 rpm until reaching less than 1 ml repeatedly for 20–30 min at room temperature (KN-70, level = 9, KUBOTA, Tokyo, Japan). On reaching 1 ml, 15 ml of phosphate-buffered saline (PBS) was added to the filter unit with the remaining supernatant and was centrifuged at 3,000 rpm until reaching less than 1 ml. This step was repeated twice using fresh PBS. Finally, the recombinant AAV vector was dispensed into each tube at an optimum volume and stored at −80°C until before starting the experiment. The recombinant AAV vector was injected with a viral titer of 4.4 × 10^13^ vg/mL for AAV9-TRE::ArchT3.0-EYFP.

### Behavioral procedures

All behavioral experiments were performed during the light cycle. Male mice were numbered and randomly assigned to an experimental group before the experiment. All behavioral experiments were performed by a researcher who was blinded to the experimental conditions. For all behavioral procedures, animals in their home cages were moved to a rack in a resting room next to the behavioral testing room and left for at least 30 min before each behavioral experiment. Different carriers were used to transfer animals from the resting room to the behavioral testing room for every behavioral test session. After the completion of all pharmacological and optogenetic experiments, the injection sites were histologically verified. Data from animals were excluded if the animals showed abnormal behavior after surgery, such as torticollis or hair pulling, or if remarkable weight loss was observed, the target area was missed, or the bilateral expression of virus was inadequate (Table S1 and S2, Excluded).

#### The context pre-exposure and immediate shock task

A previous procedure (Ohkawa et al., 2015) with minor modifications was employed. Two contexts (context A and context B) were used in this study. Context A was a cylindrical environment (diameter × height: 180 × 230 mm) with a beige floor and a wall covered with black tape. Context B was a plexiglass front and gray side- and back-walls (width × depth × height: 175 × 165 × 300 mm), and the floor consisted of stainless steel rods that were connected to a shock generator (Muromachi Kikai, Tokyo, Japan).

Mice were placed in context A (unpaired) or context B (paired) for 6 min (context pre-exposure), and then returned to their home cage. Thirty min or 1 day after context pre-exposure, the mice were given a 0.8 mA foot shock for 2 s (immediate shock, IS) in context B, 5 s after the acclimation, and returned to their home cage 1 min after the IS. After 1 day, mice were placed back into context B to test their fear memory for 3 min. Context re-exposure for 3 min was repeated as reactivation, test 1, test 2, and test 3, as shown in Figure 4 and Figure 5. At the end of each session, mice were returned to their home cages and the chambers were cleaned with 70% ethanol. All experiments were conducted using a video tracking system (Muromachi Kikai, Tokyo, Japan) to automatically measure freezing behavior. Freezing was defined as a complete absence of movement, except for respiration. We started scoring the duration of the freezing response after 1 s of sustained freezing behavior. For each test, the freezing percentage was then averaged across mice within the same group.

#### The context memory test

The context memory test was carried out as described previously (Kitamura et al., 2012). Mice were placed in context B for 6 min and were then returned to their home cages. After 1 day, mice were placed back into context B for 3 min to test their context memory, which was measured using motility. Motility was measured using a video tracking system (Muromachi Kikai, Tokyo, Japan), and was calculated as the cumulative area of movement (pixel size) per 0.1 s in the conditioning and test sessions.

#### The social interaction test

Test mice were placed into new cages prior to the experimental session and allowed to habituate to the new environment for 60 min. A 6–8 weeks old male C57BL/6J mouse that had been anesthetized with intraperitoneal injection of pentobarbital (80 mg/kg of body weight) was placed into the cage with the subject for 5 min. Social behavior was assessed by measuring the interaction time and the latency to the first interaction of test mice to anesthetized mice using a hand-held stopwatch. Interaction was defined as direct contact between the test mouse’s nose and the body of the anesthetized mouse.

### Stereotaxic virus injection

The c-fos::tTA transgenic mice were maintained with food containing 40 mg/kg Dox from weaning until the age of 12–18 weeks, at the time of surgery. The male C57BL/6J and c-fos::tTA transgenic mice were anesthetized with a pentobarbital solution (80 mg/kg of body weight; intraperitoneal injection), and the fully anesthetized mice were placed in a stereotactic apparatus (David Kopf Instruments, Tujunga, CA, USA). The male C57BL/6J and c-fos::tTA transgenic mice were injected with LV or AAV to the posterior parietal cortex (PPC), CA1, or basolateral amygdala (BLA). The positions (in mm) were as follows: (i) −2.0 AP, ±1.5 ML from bregma, −0.5 DV from the brain surface for the PPC; (ii) −2.0 AP, ±1.5 ML, −1.5 DV from bregma for the CA1; (iii) −1.5 AP, ±3.3 ML, - 4.9 DV from bregma for the BLA. Borosilicate glass capillaries (G100T-4, outer diameter: 1.0 mm; inner diameter: 0.78 mm; Warner Instruments, Hamden, CT) were pulled using a Micropipette Puller (P-1000, Sutter Instrument, Novato, CA, USA; heat: 778, pull: 25, vel: 150, time: 250, and pressure: 500). The tip of the glass capillary was trimmed with scissors under an operating microscope to have an outer diameter of about 50 μm. For the virus delivery, 1 μl LV (delivery rate, 0.1 μl/min) or 0.3 μl AAV (delivery rate, 0.06 μl/min) was injected using a motorized stereotaxic injector (Narishige, Tokyo, Japan) and a Hamilton microsyringe coupled with a glass capillary. After the injection, the capillary was left in place for 10 min and then slowly withdrawn. To insert an optical fiber into the PPC, CA1, BLA, or ACC, guide cannulas (C316GS-4/SPC, Plastics One, Roanoke, VA, USA) were implanted slightly above the target coordinates of each region. The coordinate for the ACC (in mm) was +1.0 AP, +0.5 ML from bregma, −0.5 DV from the brain surface. The guide cannulas were fixed on the skull using dental cement. A dummy cannula with a cap was used to protect the guide cannula from clogging up. After recovery from anesthesia in a stable-temperature incubator, mice were returned to their home cages.

### Optic fiber placement and optical stimulation

Optical fiber unit placement was carried out as described previously (Nomoto et al., 2016; Yokose et al., 2017). For the placement of the optical fiber units, mice were anesthetized with 2.0% isoflurane and the dummy cannulas were removed from the guide cannulas. The optical fiber unit, composed of a plastic cannula body, was a two-branch-type unit with an optic fiber diameter of 0.250 mm (COME2-DF2-250, Lucir, Tsukuba, Japan). The optical fiber unit was inserted into the guide cannula, and the bodies of the guide cannula and the optical fiber unit were tightly connected using an adhesive agent (Kokuyo, Osaka, Japan). The tips of the optical fibers were targeted slightly above the target coordinates of each region. The fiber unit connected with the mouse was attached to an optical swivel (COME2-UFC, Lucir, Tsukuba, Japan), which was connected to a laser (200 mW, 473 nm and 589 nm, COME-LB473/589/200, Lucir, Tsukuba, Japan) via a main optical fiber. The delivery of light pulses was controlled by a schedule stimulator (COME2-SPG-2, Lucir, Tsukuba, Japan) in time-lapse mode. For the silencing of cell assemblies by ArchT or NpHR, optical illumination (continuous 589 nm light, 12-14 mW at the fiber tip) was delivered, as shown in each figure, except that optical illumination in the IS session was delivered 1 min before starting. For the activation of cell assemblies by ChR2, optical illumination (473 nm light, 20 Hz, 10 ms, 6-7 mW at the fiber tip) was delivered. Two hours after the optical illumination had ended, mice were anesthetized with 2.0% isoflurane, the optic fiber unit was detached, and the mice were returned to their home cage. All mice were sacrificed and immunostained to verify the coordinates after behavioral experiments. Mice in which no opsin expression was confirmed were excluded from each experiment (Table S1 and S2, Excluded).

### Drug infusion

Mice anesthetized with pentobarbital (80 mg/kg of body weight) were implanted bilaterally with stainless steel guide cannulas (C316GS-4/SPC, Plastics One, Roanoke, VA, USA) using the following stereotactic coordinates (in mm): (i) −2.0 AP, ±1.5 ML from bregma, −0.3 DV from the brain surface for injection into the PPC; (ii) −2.0 AP, ±1.5 ML, −0.5 DV from bregma for the CA1 region; (iii) −1.5 AP, ±3.3 ML, −3.4 DV from bregma for the BLA. After surgery, a dummy cannula was inserted into the guide cannula, and mice were allowed to recover for at least 14 days in individual home cages. To prevent disruption of the target region, the injection cannula (C316IS-4/SPC, Plastics One, Roanoke, VA, USA) extended beyond the tip end of the guide cannula by 0.2 mm for the PPC, and 1.0 mm for the CA1 and the BLA. A sodium channel blocker (lidocaine-hydrochloride; Sigma, St. Louis, MO, USA, L5647) was used to inactivate the PPC, CA1 region, or BLA. Mice were briefly anesthetized with isoflurane to facilitate the insertion of the injection cannula. Lidocaine-hydrochloride (4%, 1 μl) or PBS was infused into each brain region at a rate of 0.333 μl/min, which was controlled by a microsyringe pump (Legato111, KD Scientific Inc., Holliston, MA, USA). After infusion, the injection cannula was left in place for 3 min to allow for drug diffusion. After behavioral experiments, all mice were injected 0.5 μl of 0.5 μM Rhodamine B (Sigma, St. Louis, MO, USA) through the guide cannula to verify the coordinates. Mice in which no Rhodamine B expression was confirmed were excluded from each experiment (Table S2, Excluded).

### Tracer injection

Biotin Dextran Amine (BDA, Thermo Fisher Scientific, Waltham, MA, USA, D1817) and fluorescent retrograde tracer cholera toxin B subunit conjugated to fluorophore Alexa 488 (CTB488, Thermo Fisher Scientific, Waltham, MA, USA, C34775) were used to label neurons that projected from the target regions. BDA and CTB488 were unilaterally delivered into the PPC and ACC of male C57BL6J mice using a glass micropipette at the following coordinates (in mm): (i) −2.0 AP, +1.5 ML from bregma, −0.5 DV from the brain surface for the PPC; (ii) +1.0 AP, +0.5 ML from bregma, −0.5 DV from the brain surface for the ACC. Mice were injected with 200 nl of 10% BDA in the PPC or 200 nl of 1% CTB488 in the ACC, and mice were perfused 7 days or 5 days after injection. BDA was visualized by staining with streptavidin-Alexa Fluor 555 (1:1000, Life Technologies, Carlsbad, CA, USA).

### Fluorescent in situ hybridization

Five minutes after the behavioral experiments, brains were rapidly extracted, frozen, and stored at −80°C before sectioning. Twenty μm-thick coronal sections of mouse brain were cut using a cryostat, air-dried, and mounted onto slides.

Fluorescein-labeled arc riboprobes were synthesized using a transcription kit (MaxiScript, Ambion Inc., Austin, TX, USA) and premixed RNA labeling nucleotide mixes containing fluorescein-labeled UTP (Roche Diagnostics, Indianapolis, IN, USA). Riboprobes were purified using G-50 spin columns (GE Healthcare, Waukesha, WI, USA). The plasmid containing nearly full-length cDNA of the arc transcript was used to generate probes.

Fluorescent in situ hybridization (FISH) was carried out as described previously (Nomoto et al., 2016; Ohkawa et al., 2015; Yokose et al., 2017). For Arc CatFISH, brain sections were fixed in fresh 4% paraformaldehyde/0.1 M phosphate buffer at room temperature. All steps were performed at room temperature unless otherwise indicated. Sections were acetylated with 0.5% acetic anhydride/1.5% triethanolamine, dehydrated using 1:1 methanol/acetone solutions, and equilibrated in 2× saline sodium citrate buffer (SSC). Sections were then prehybridized for 30 min in prehybridization buffer [50% formamide, 5% sheared salmon sperm DNA (Sigma, St. Louis, MO, USA), 0.3% yeast tRNA (Sigma, St. Louis, MO, USA), 5× SSC, 5× Denhart’s solution (Sigma, St. Louis, MO, USA)]. The antisense riboprobe was diluted in hybridization buffer at a final concentration of 1.5 μg/ml and applied to each slide. Coverslips were added to the slides, and hybridization was carried out at 56°C for 16 hours. Sections were dipped in 2× SSC and rinsed in 0.5× SSC followed by 0.5× SSC at 56°C, which included an earlier wash step in 2× SSC with RNase A (10 μg/ml) at 37°C. Sections were again washed in 0.5× SSC and endogenous peroxidase activity was inhibited by incubation in 2% H_2_O_2_ in 1× SSC for 15 min. Sections were blocked with TSA blocking reagent (PerkinElmer, Waltham, MA, USA) and incubated with an anti-fluorescein horseradish peroxidase conjugate (1:500, PerkinElmer, Waltham, MA, USA) overnight at 4°C. Sections were washed three times in Tris-buffered saline (with 0.05% Tween-20), and the conjugate was detected using the Fluorescein TSA Plus System (1:10, PerkinElmer, Waltham, MA, USA) for 2 h. Sections were washed in PBS and the nuclei were counterstained with DAPI(1 μg/ml, Roche Diagnostics, Indianapolis, IN, USA, 10236276001) for 20 min. Finally, the sections were washed with PBS and mounted with Permafluor mounting medium (Lab Vision Corporation, Fremont, CA, USA).

Images were acquired using a Zeiss LSM 780 confocal microscope with a 40×/1.2 NA objective lens. PMT assignments, pinhole sizes, and contrast values were kept constant. Images of the CA1 region and BLA were acquired by collecting z-stacks (1 μm-thick optical sections). Using the Zen software, each cell was characterized through several serial sections, and only cells containing whole nuclei were included in the analysis. The details of the classification analysis have been described before (Guzowski et al., 1999). Small, bright, uniformly DAPI-stained nuclei (from putative glial cells) were not analyzed. All other whole nuclei were analyzed from top to bottom. The designation ‘cytoplasm-positive (Cyto) neurons’ was given to cells containing perinuclear/cytoplasmic labeling in multiple optical sections. The designation ‘nuclear-positive (Nuc) neurons’ was given to cells containing two small, intense intranuclear fluorescent foci, and the designation ‘Nuc and Cyto double positive (Cyto/Nuc)’ was given to cells containing both intranuclear and cytoplasmic Arc-positive signals. Three sections corresponding to each region of interest (ROI) were chosen from each mouse.

### Immunohistochemistry

Mice were deeply anesthetized with an overdose of pentobarbital solution and perfused transcardially with PBS, pH 7.4, followed by 4% paraformaldehyde in PBS. The brains were removed and further post-fixed by immersion in 4% paraformaldehyde in PBS for 24 h at 4°C. Each brain was equilibrated in 30% sucrose in PBS, and then frozen in dry ice powder. For EYFP and c-Fos staining, coronal sections were cut on a cryostat at a thickness of 50 μm, 1.2 mm to 0.8 mm, and −1.4 to −1.8 mm from bregma for the ACC and BLA, respectively, and transferred to 12-well cell culture plates (Corning, Corning, NY, USA) containing PBS. After washing with PBS, the floating sections were treated with blocking buffer, 5% normal donkey serum (S30, Chemicon by EMD Millipore, Billerica, MA, USA) in PBS with 0.2% Triton X-100 (PBST), at room temperature for 1h. Reactions with primary antibodies were performed in blocking buffer containing rabbit anti-GFP (1:1000, Invitrogen, Carlsbad, CA, USA, A11122) and/or goat anti-c-Fos (1:1000, SantaCruz, Santa Cruz, CA, USA, SC-52G) antibodies for double staining of EYFP and c-Fos, chicken anti-GFP (1:1000, abcam, Cambridge, UK, ab13970), and/or rabbit anti-Egr-1 (1:1000, SantaCruz, Santa Cruz, CA, USA, SC-189) antibodies for double staining of EYFP and Zif268 at 4°C for 1 or 2 days. After three 10 min washes with PBS, the sections were incubated with donkey anti-rabbit IgG-Alexa Fluor 488 (1:300, Life Technologies, Carlsbad, CA, USA, A21206) and/or donkey anti-goat IgG-Alexa Fluor 546 (1:300, Life Technologies, Carlsbad, CA, USA, A11056) secondary antibodies for double staining of EYFP and c-Fos, and donkey anti-chicken IgG-Alexa Fluor 488 (1:300, Jackson ImmunoResearch Laboratories, Inc, West Grove, PA, USA, 703-545-155) and donkey anti-rabbit IgG-Alexa Fluor 546 (1:300, Life Technologies, Carlsbad, CA, USA, A10040) secondary antibodies for double staining of EYFP and Zif268 at room temperature for 3 h. Sections were treated with DAPI (1 μg/ml, Roche Diagnostics, Indianapolis, IN, USA, 10236276001) and then washed with PBS three times, 10 min/wash. Mounting of sections on glass slides was performed with ProLong Gold antifade reagents (Invitrogen, Carlsbad, CA, USA). Images were acquired using a Zeiss LSM 780 confocal microscope with a 20×/1.2 NA objective lens. PMT assignments, pinhole sizes, and contrast values were kept constant. For quantification of the number of positive cells, images of the ROI were acquired by collecting z-stacks (1 μm-thick optical sections). Quantification of the number of c-Fos-positive or Zif268-positive cells in the ROI was performed by an experimenter who was blinded to the experimental conditions.

### Statistical analyses

Statistical analysis was performed using GraphPad Prism version 6 (GraphPad Software, San Diego, CA, USA). All data are presented as the mean ± SEM. The number of animals used is indicated by “n”. Comparisons between two groups were made using unpaired Student’s t-tests. Multiple group comparisons were assessed using a one-way, two-way, or repeated measures analysis of variance (ANOVA), followed by the Tukey–Kramer test when significant main effects or interactions were detected. The null hypothesis was rejected at the P < 0.05 level. All statistical data are described in Table S1 and S2.

## Notes

### Competing Interest Statement

The authors have declared no competing interest.

