## Supplementary Figures for "Cortical cell ensemble control of past experience-dependent memory updating"

### Supplementary Figure

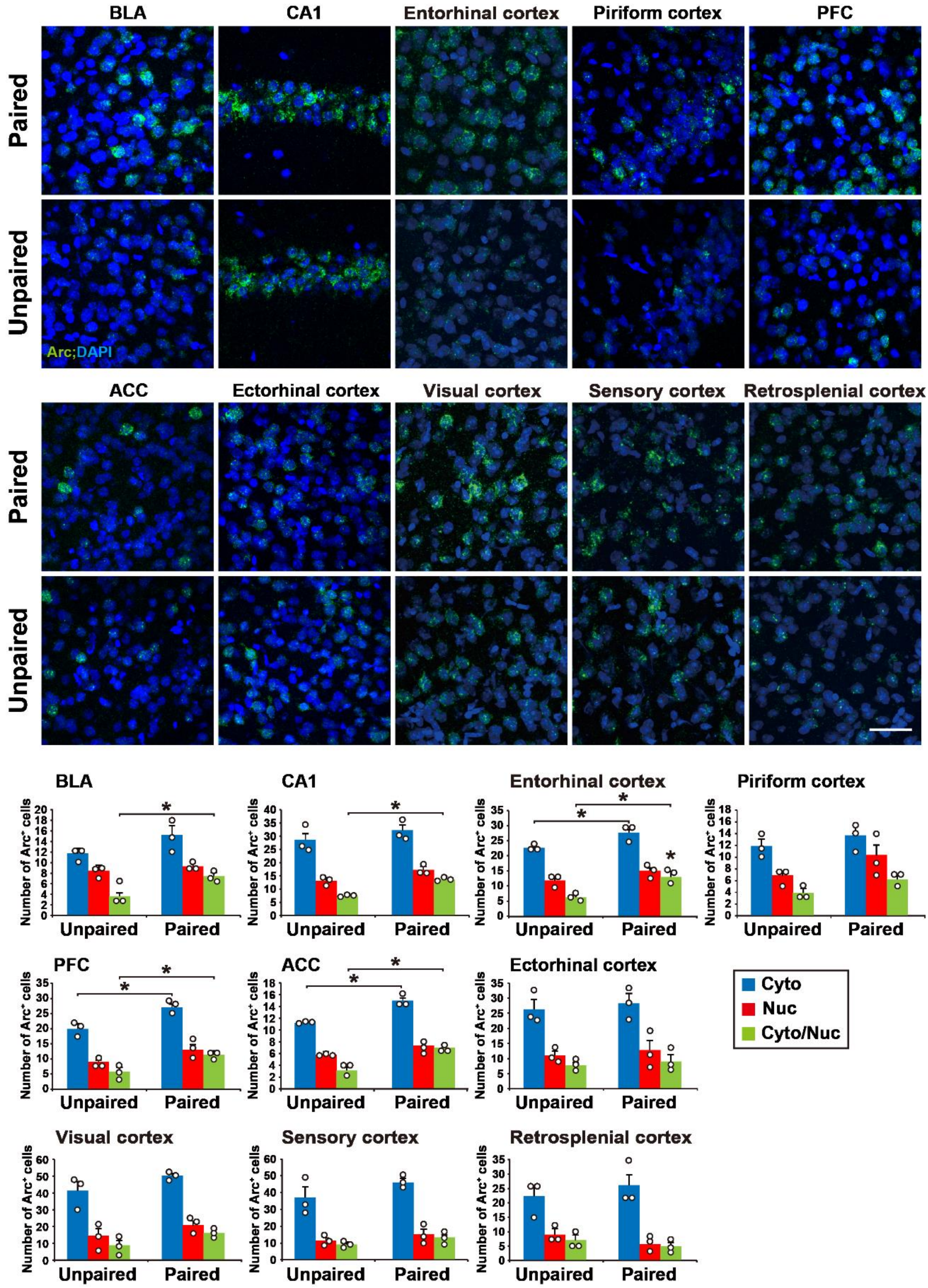

**Figure S1. Multiple brain regions respond to the pre-exposed context and IS, Related to Figure 1**

Representative images of the Arc CatFISH analysis in several brain regions (top). The Arc RNA signal and DAPI nuclear staining are shown in green and blue, respectively. Scale bar, 50 μm. Number of Arc+ cells (bottom) (n = 3 mice/group).

Error bars indicate the mean ± s.e.m. *$P < 0.05$. For details of statistical data, see Table S2. Cyto, cytoplasmic; Nuc, nuclear.

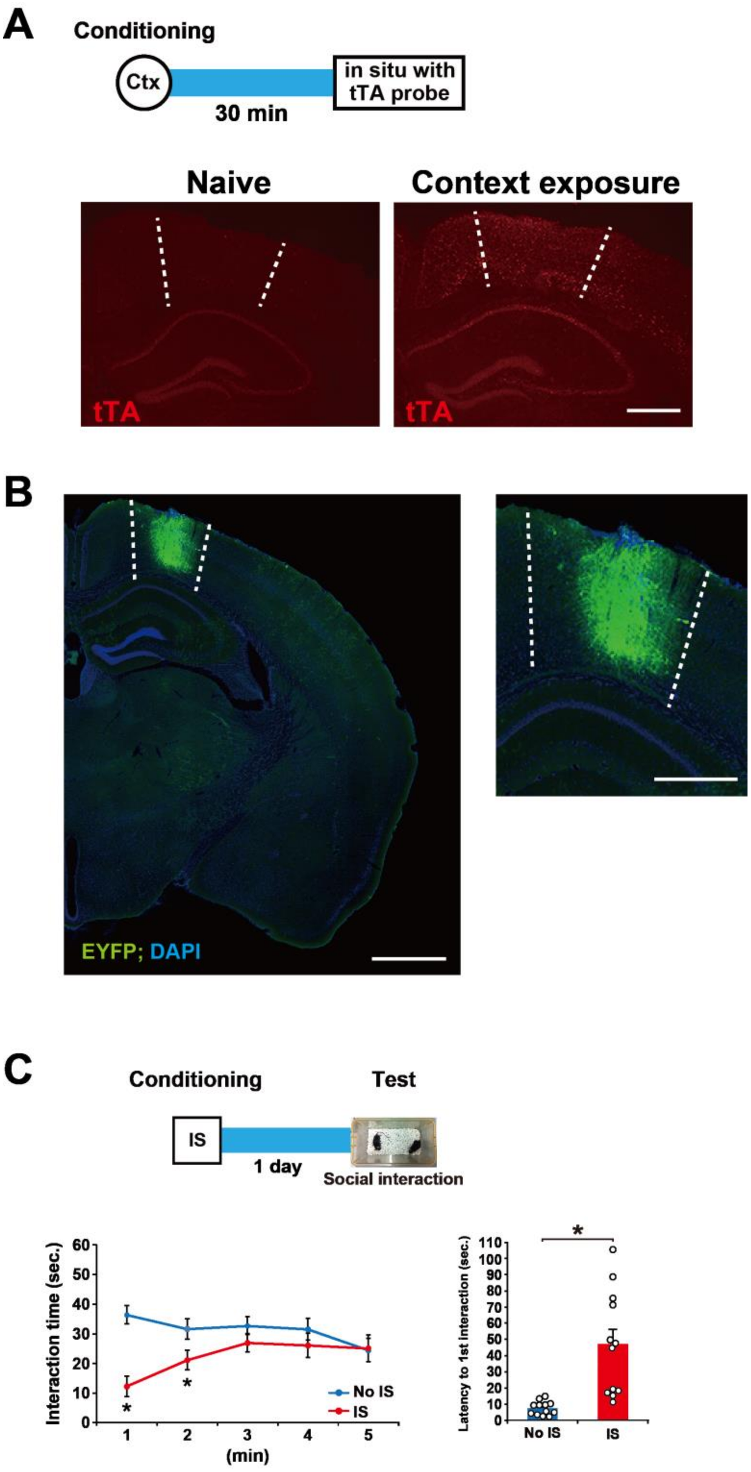


**Figure S2. Histological and behavioral studies, Related to Figure 2**

(A) tTA mRNA is expressed in the PPC. Schema of the behavioral and in situ experiments. Mice were sacrificed 30 min after context exposure (top). Control mice without context exposure were used as naïve mice. Representative images of the tTA in situ analysis in naïve and context-exposed mice (bottom). Red dots represent the tTA mRNA signal. Broken lines indicate the PPC boundaries. Scale bar, 500 μm. Ctx, context.

(B) Expression of ArchT-EYFP in the PPC. Representative labeling pattern of PPC cells with ArchT-EYFP protein in a c-fos::tTA transgenic mice that was conditioned with context exposure at 2 days after OFF Dox (left). The signal of fluorescent protein and DAPI nuclear staining at 1 day after the conditioning are shown in green and blue, respectively. This section was taken from bregma -1.94 mm. Scale bar, 1 mm. An enlarged image of the injection area (right). Broken lines indicate the PPC boundaries. Scale bar, 500 μm.

(C) IS affects the social interaction. Schematic of the behavioral experiment (top). Graph shows the interaction time (bottom left) and the latency to the first interaction (bottom right) during the test (bottom) (No-IS, n = 12 mice; IS, n = 12 mice).

Error bars indicate the mean ± s.e.m. *P, < 0.05. IS, immediate shock. For details of statistical data, see Table S2.

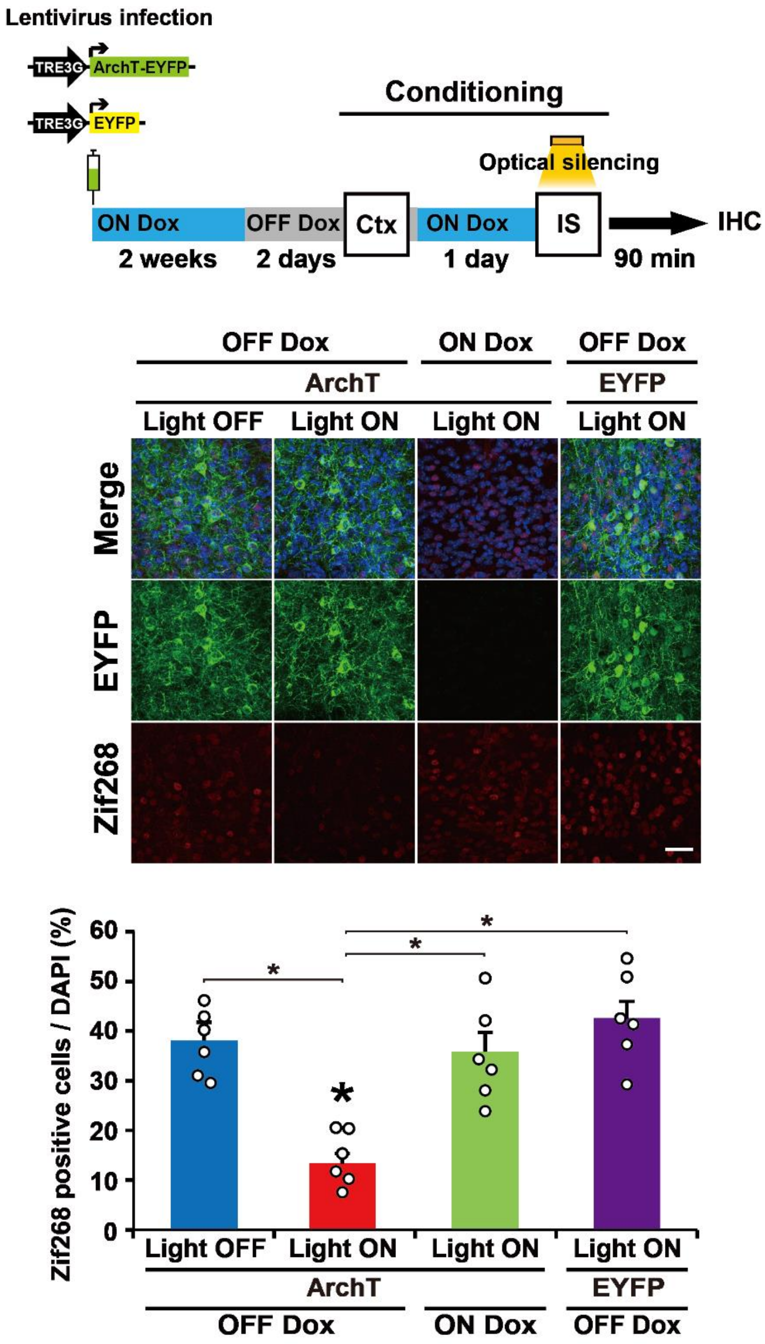


**Figure S3. ArchT-EYFP functions in the Light ON and OFF Dox condition, Related to Figure 2**

The behavioral experiment with optical silencing (top). Blue and gray bars indicate the presence or absence of Dox, respectively. A representative image of Zif268 expression 90 min after an IS session with or without optical silencing (middle). Scale bar, 100 μm.

The proportion of Zif268-positive cells in each group (bottom) (n = 6 sections from 2 mice/group). Error bars indicate the mean ± s.e.m. *$P < 0.05$. For details of statistical data, see Table S2.

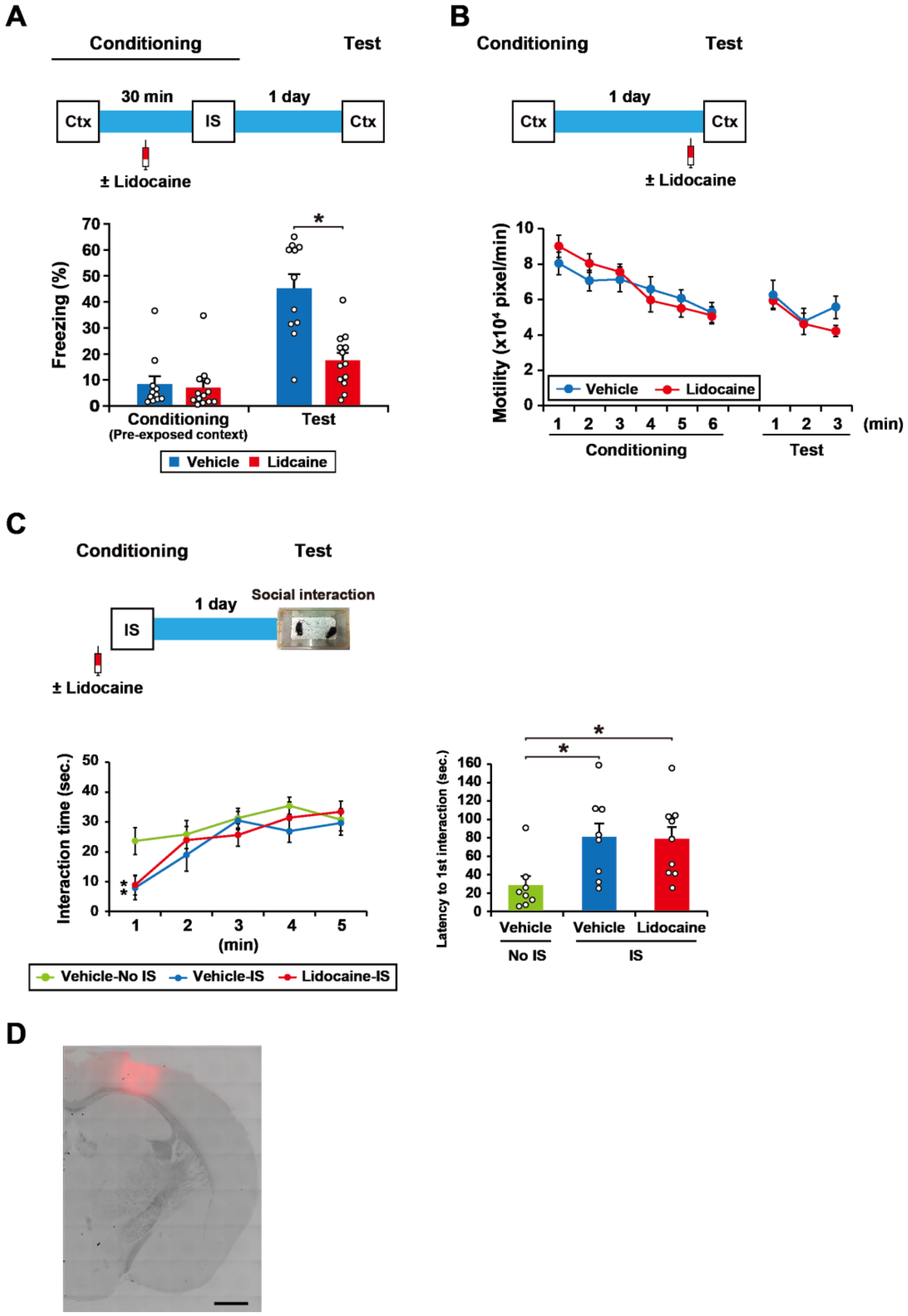


**Figure S4. Lidocaine injection in the PPC blocks memory association, Related to Figure 2**

(A) Schematic of the behavioral experiment (top). The mice were pre-exposed to context B for 6 min, and then received IS in the same context at intervals of 30 min. The sodium channel blocker lidocaine was injected into the PPC 15 min before IS exposure. The effect of the lidocaine injection on memory association is shown (bottom). The graph shows the freezing level during the conditioning and test session (vehicle, n = 11 mice, lidocaine, n = 13 mice).

(B) Schematic of the behavioral experiment (top). The graph shows motility at the conditioning and test session (bottom) (vehicle, n = 7 mice, lidocaine, n = 8 mice).

(C) Schematic of the behavioral experiment (top). The graph shows the interaction time (left) and the latency to the first interaction (right) during the test session (bottom) (vehicle-No IS, n = 8 mice, vehicle-IS, n = 8 mice, lidocaine-IS (+), n = 9 mice). Error bars indicate the mean ± s.e.m. *$P < 0.05$.

(D) Representative image of a rhodamine injection into the PPC. All mice were injected with rhodamine in the PPC through a guide cannula to check the coordinates after the behavioral experiments. Scale bar, 1 mm.

Ctx, context; IS, immediate shock. For details of statistical data, see Table S2.

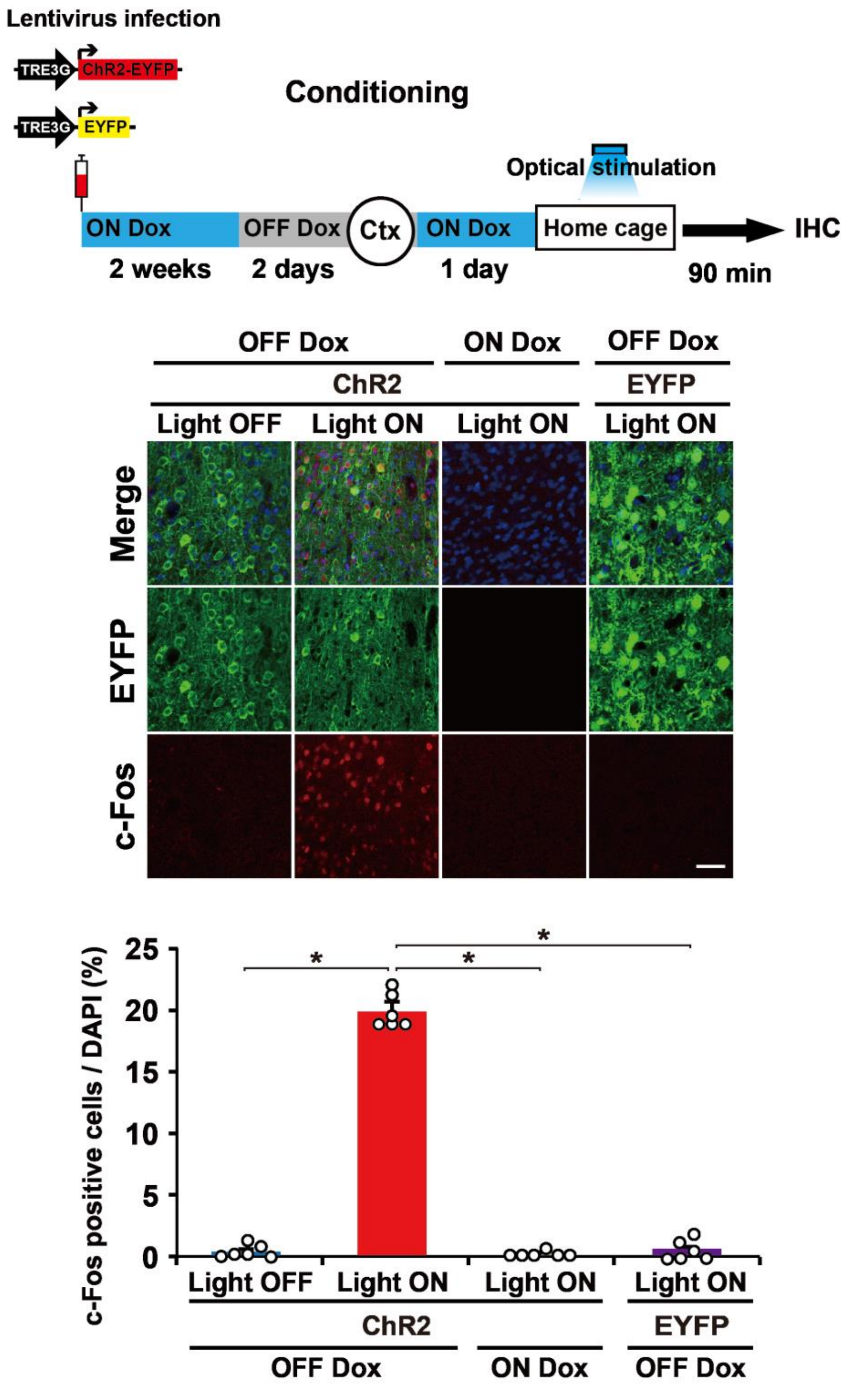


**Figure S5. ChR2-EYFP functions in the Light ON and OFF Dox condition, Related to Figure 3**

The behavioral experiment with optical stimulation (top). Blue and gray bars indicate the presence or absence of Dox, respectively. Mice were sacrificed 90 min after optical stimulation. A representative image of c-Fos expression (middle). Scale bar, 100 μm. The proportion of c-Fos-positive cells in each group (bottom) (n = 6 sections from 2 mice/group). Error bars indicate the mean ± s.e.m. *P < 0.05. For details of statistical data, see Table S2.

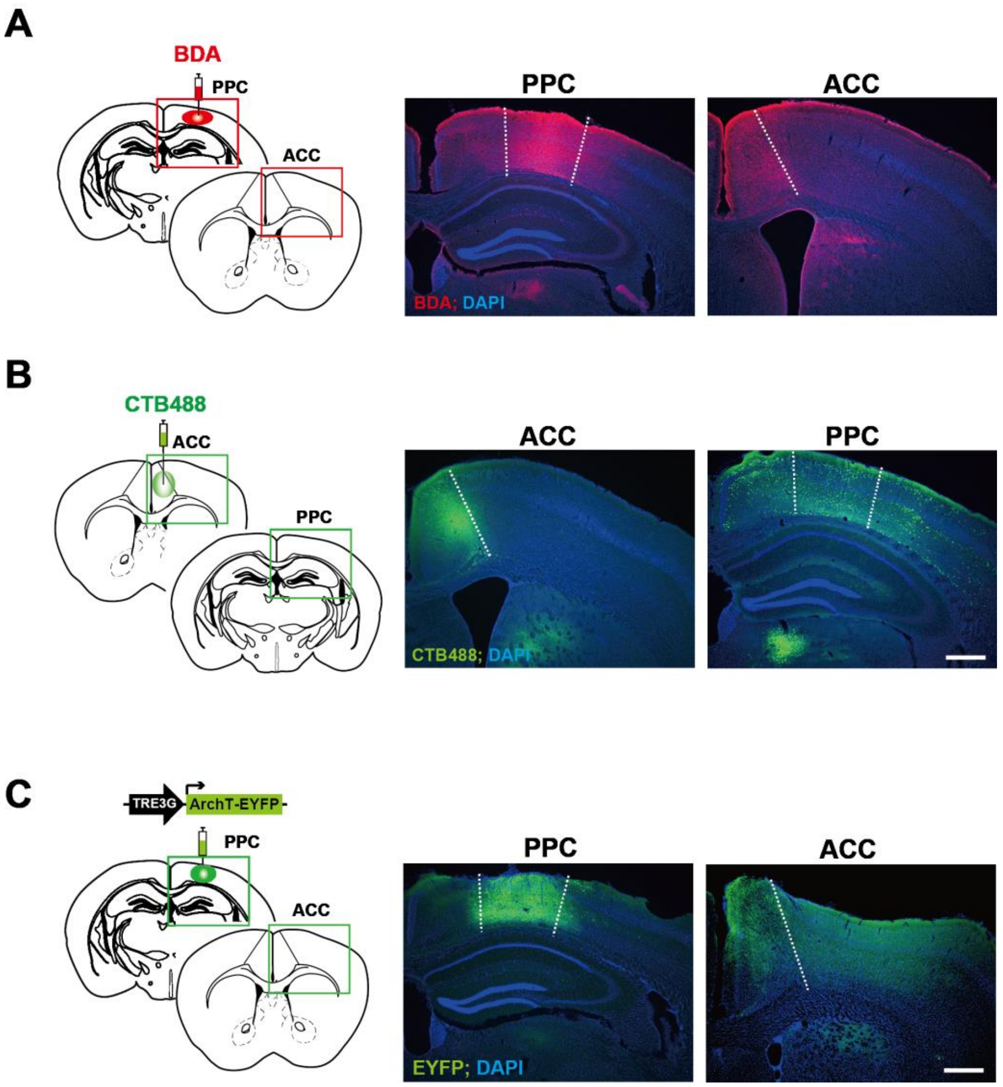


**Figure S6. Histological studies, Related to Figure 7**

(A and B) PPC neurons project to the ACC. Schematics showing the BDA injection area and observational area (panel A left). BDA was injected into the PPC. Magnified images of the red square areas are shown on the right. Representative images of brain sections of the PPC and ACC immunostained with the streptavidin-Alexa Fluor 555 (red, panel A right). Schematics showing the CTB488 injection area and observational area (panel B left). CTB488 was injected into the ACC. Magnified images of the green square areas are shown on the right. Representative images of ACC and PPC brain sections are shown (panel B right). Scale bar, 500 μm.

(C) Expression of ArchT-EYFP in neurons in the PPC and PPC axons in the ACC. Schematics showing labeling of the PPC neurons in c-fos::tTA transgenic mice with the AAV9 TRE3G::ArchT-EYFP (left). Representative images of PPC and ACC brain sections are shown (right). Scale bar, 500 μm.

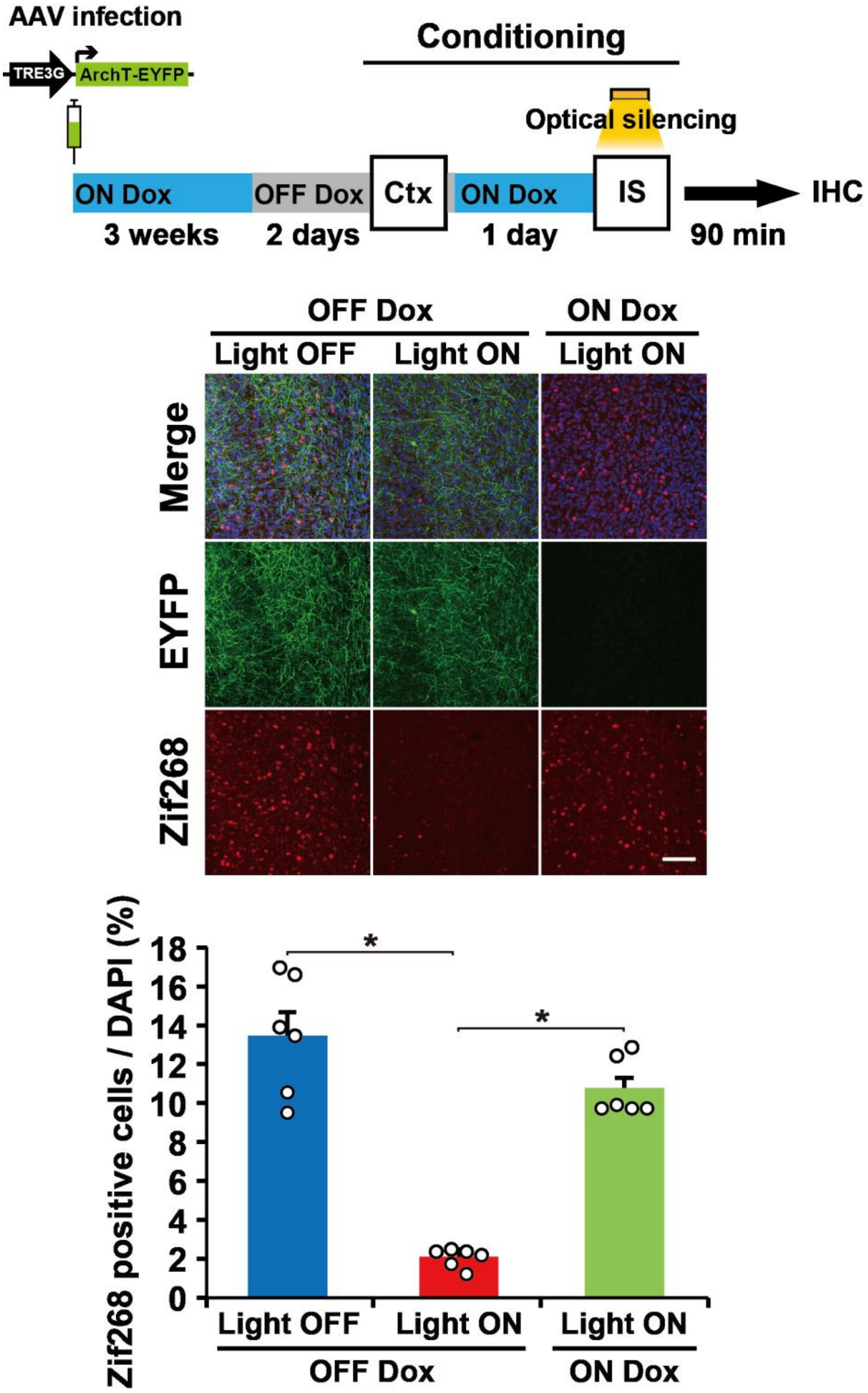


**Figure S7. ArchT-EYFP in ACC functions in the Light ON and OFF Dox condition, Related to Figure 7**

The behavioral experiment with optical silencing (top). Blue and gray bars indicate the presence or absence of Dox, respectively. Optical silencing to the ACC was delivered during an IS session of conditioning. A representative image of Zif268 expression 90 min after the IS session with or without optical silencing (middle). Scale bar, 100 μm. The proportion of Zif268-positive cells in each group (bottom) (n = 6 sections from 2 mice/group). Error bars indicate the mean ± s.e.m. *$P < 0.05$. For details of statistical data, see Table S2.
