## Supplementary Table1 for "Cortical cell ensemble control of past experience-dependent memory updating"

**Table S1. Sampling and statistical analysis, Related to Figure 1 - Figure 7.**

| Fig. # | Category | Group | Sample size (n) | | Score | Statistical test | Degree of Freedom & F/t Value | p-value | Significance? |
|---|---|---|---|---|---|---|---|---|---|
| | | | Exact size (n) | Excluded (n) | | | | | |
| 1B | Freezing (%) in Conditioning | PEC | 9 | 0 | 3.6 ± 1.22% | | | | |
| | | IS | 9 | 0 | | Unpaired t test | t16 = 0.2515 | P = 0.8047 | N.S. |
| | | Paired | 9 | 0 | 3.9 ± 1.13% | | | | |
| | Freezing (%) in Test | PEC | 9 | 0 | 4.6 ± 1.14% | | | | |
| | | IS | 9 | 0 | 8.8 ± 2.45% | One-way ANOVA | F2,24 = 59.96 | P < 0.0001 | **** |
| | | Paired | 9 | 0 | 33.1 ± 4.68% | | | | |
| 1C | Freezing (%) | 30 min interval Unpaired | 8 | 0 | Conditioning; 16.4 ± 2.26%, Test; 16.1 ± 3.79% | Two-way ANOVA with RM | session; F1,14 = 40.44<br>group; F1,14 = 6.596<br>session x group; F1,14 = 41.58 | session; P < 0.0001<br>group; P = 0.0223<br>session x group; P < 0.0001 | session; ****<br>group; *<br>session x group; **** |
| | | 30 min interval Paired | 8 | 0 | Conditioning; 10.9 ± 2.99%, Test; 45.0 ± 5.23% | | | | |
| | | 1 day interval Unpaired | 8 | 0 | Conditioning; 12.1 ± 3.21%, Test; 13.9 ± 3.80% | Two-way ANOVA with RM | session; F1,14 = 22.47<br>group; F1,14 = 6.741<br>session x group; F1,14 = 17.95 | session; P = 0.0003<br>group; P = 0.0211<br>session x group; P = 0.0008 | session; ***<br>group; *<br>session x group; *** |
| | | 1 day interval Paired | 8 | 0 | Conditioning; 8.9 ± 3.57%, Test; 41.2 ± 5.63% | | | | |
| 1F | Arc+ cells / DAPI (%) | Unpaired | 6 | 0 | Cyto; 28.9 ± 3.24%, Nuc; 5.5 ± 0.57%, Cyto/Nuc; 3.5 ± 0.24% | One-way ANOVA | Cytoplasmic arc; F2,12 = 3.837<br>Nuclear arc; F2,12 = 20.38<br>Cytoplasmic and nuclear arc; F2,12 = 26.95 | Cytoplasmic arc; P = 0.0515<br>Nuclear arc; P = 0.0001<br>Cytoplasmic and nuclear arc; P < 0.0001 | Cytoplasmic arc; N.S.<br>Nuclear arc; ***<br>Cytoplasmic and nuclear arc; **** |
| | | Paired | 6 | 0 | Cyto; 39.3 ± 2.11%, Nuc; 12.6 ± 0.96%, Cyto/Nuc; 10.5 ± 0.98% | Unpaired t test (for vs Chance) | Unpaired; t10= 4.462<br>Paired; t10= 4.796<br>No IS; t4 = 3.344 | Unpaired; P= 0.0012<br>Paired; P= 0.0007<br>No IS; P= 0.0287 | Unpaired; **<br>Paired; ***<br>No IS; * |
| | | No IS | 3 | 0 | Cyto; 35.6 ± 3.36%, Nuc; 7.5 ± 1.21%, Cyto/Nuc; 5.6 ± 0.78% | | | | |
| 2B | Freezing (%) | ArchT-Light OFF | 9 | 0 | Conditioning; 8.1 ± 1.74%, Test; 44.9 ± 4.01% | Two-way ANOVA with RM | session; F1,29 = 147.1<br>group; F2,29 = 5.579<br>session x group; F2,29 = 7.879 | session; P < 0.0001<br>group; P = 0.0089<br>session x group; P = 0.0018 | session; ****<br>group; **<br>session x group; ** |
| | | ArchT-Light ON | 11 | 0 | Conditioning; 9.5 ± 2.23%, Test; 24.6 ± 3.99% | | | | |
| | | EYFP-Light ON | 12 | 0 | Conditioning; 12.0 ± 2.69%, Test; 41.3 ± 2.90% | | | | |
| 2C | Freezing (%) | Circle labeled | 12 | 0 | Conditioning (Circle-context); 4.7 ± 0.66%, Conditioning (Square-context); 7.9 ± 1.59%, Test; 44.1 ± 4.81% | Two-way ANOVA with RM | session; F2,44 = 131.0<br>group; F1,22 = 14.78<br>session x group; F2,44 = 16.64 | session; P < 0.0001<br>group; P = 0.0009<br>session x group; P < 0.0001 | session; ****<br>group; ***<br>session x group; **** |
| | | Square labeled | 12 | 0 | Conditioning Circle-context; 5.5 ± 0.97%, Conditioning Square-context; 6.0 ± 0.99%, Test; 23.8 ± 2.30% | | | | |
| 2D | Motility (pixel/min) Target: PPC | ArchT-Light OFF | 10 | 0 | Conditioning;<br>1 min; 86846 ± 5272 pixel, 2 min; 70844 ± 6725 pixel, 3 min; 56555 ± 4866 pixel,<br>4 min; 45718 ± 3409 pixel, 5 min; 45532 ± 2982 pixel, 6 min; 46405 ± 4473 pixel,<br><br>Test;<br>1 min; 33327 ± 5531 pixel, 2 min; 26145 ± 3957 pixel, 3 min; 33446 ± 4277 pixel | Two-way ANOVA with RM | Conditioning Session;<br>time; F5,144 = 34.44<br>group; F2,144 = 1.168<br>time x group; F10,144 = 0.4690<br><br>Test session;<br>time; F2,72 = 5.013<br>group; F2,72 = 0.841<br>time x group; F4,72 = 0.7140 | Conditioning Session;<br>time; P < 0.0001<br>group; P = 0.3140<br>time x group; P = 0.9077<br><br>Test session;<br>time; P = 0.0092<br>group; P = 0.4356<br>time x group; P = 0.5851 | Conditioning Session;<br>time; ****<br>group; N.S.<br>time x group; N.S.<br><br>Test session;<br>time; **<br>group; N.S.<br>time x group; N.S. |
| | | ArchT-Light ON | 10 | 0 | Conditioning;<br>1 min; 95929 ± 8908 pixel, 2 min; 70113 ± 6550 pixel, 3 min; 52386 ± 5710 pixel,<br>4 min; 41445 ± 3653 pixel, 5 min; 41623 ± 3577 pixel, 6 min; 38405 ± 2741 pixel,<br><br>Test;<br>1 min; 42736 ± 3870 pixel, 2 min; 29996 ± 4097 pixel, 3 min; 30547 ± 4790 pixel | | | | |
| | | EYFP-Light ON | 7 | 0 | Conditioning;<br>1 min; 92270 ± 6524 pixel, 2 min; 70335 ± 6794 pixel, 3 min; 59980 ± 7856 pixel,<br>4 min; 54518 ± 6741 pixel, 5 min; 47799 ± 3695 pixel, 6 min; 45413 ± 4996 pixel,<br><br>Test;<br>1 min; 39571 ± 6976 pixel, 2 min; 23830 ± 5477 pixel, 3 min; 25186 ± 2771 pixel | | | | |
| | Motility (pixel/min) Target: CA1 | ArchT-Light OFF | 12 | 0 | Conditioning;<br>1 min; 87004 ± 5881 pixel, 2 min; 71926 ± 5358 pixel, 3 min; 69915 ± 4195 pixel,<br>4 min; 59864 ± 3278 pixel, 5 min; 54537 ± 2559 pixel, 6 min; 46741 ± 2672 pixel<br><br>Test;<br>1 min; 36302 ± 4323 pixel, 2 min; 29475 ± 5473 pixel, 3 min; 32384 ± 4696 pixel | Two-way ANOVA with RM | Conditioning session;<br>time; F5,204 = 27.87<br>group; F2,204 = 0.9729<br>time x group; F10,204 = 0.4125<br><br>Test session;<br>time; F2,102 = 1.457,<br>group; F2,102 = 20.62,<br>time x group; F4,102 = 0.3644 | Conditioning session;<br>time; P < 0.0001<br>group; P = 0.3797<br>time x group; P = 0.9397<br><br>Test session;<br>time; P = 0.2378<br>group; P < 0.0001<br>time x group; P = 0.8334 | Conditioning session;<br>time; ****<br>group; N.S.<br>time x group; N.S.<br><br>Test session;<br>time; N.S.<br>group; ****<br>time x group; N.S. |
| | | ArchT-Light ON | 12 | 0 | Conditioning;<br>1 min; 86810 ± 4991 pixel, 2 min; 67213 ± 5479 pixel, 3 min; 66419 ± 4554 pixel,<br>4 min; 48235 ± 5903 pixel, 5 min; 55877 ± 5208 pixel, 6 min; 47262 ± 6633 pixel<br><br>Test;<br>1 min; 66052 ± 5315 pixel, 2 min; 59284 ± 6265 pixel, 3 min; 52966 ± 7283 pixel | | | | |
| | | EYFP-Light ON | 13 | 0 | Conditioning;<br>1 min; 90508 ± 3722 pixel, 2 min; 68503 ± 4718 pixel, 3 min; 74264 ± 4500 pixel,<br>4 min; 55114 ± 5655 pixel, 5 min; 53087 ± 4934 pixel, 6 min; 51912 ± 3699 pixel<br><br>Test;<br>1 min; 40207 ± 5468 pixel, 2 min; 34212 ± 4692 pixel, 3 min; 37770 ± 4492 pixel | | | | |
| 2E | Interaction time (sec.) Target: PPC | ArchT-Light OFF | 10 | 0 | 1 min: 5.7 ± 3.23 sec, 2 min; 24.3 ± 4.95 sec, 3 min; 24.1 ± 3.97 sec,<br>4 min; 29.7 ± 5.36 sec, 5 min; 29.3 ± 6.78 sec | Two-way ANOVA with RM | time; F4,120 = 11.52<br>group; F2,120 = 0.5590<br>time x group; F8,120 = 0.4154 | time; P < 0.0001<br>group; P = 0.5733<br>time x group; P = 0.9098 | time; ****<br>group; N.S.<br>time x group; N.S. |
| | | ArchT-Light ON | 10 | 0 | 1 min: 10.2 ± 3.81 sec, 2 min; 21.1 ± 5.86 sec, 3 min; 26.8 ± 6.65 sec,<br>4 min; 33.6 ± 4.07 sec, 5 min; 35.8 ± 4.03 sec | | | | |
| | | EYFP-Light ON | 7 | 0 | 1 min: 5.3 ± 4.58 sec, 2 min; 16.3 ± 5.41 sec, 3 min; 30.3 ± 4.14 sec,<br>4 min; 34.3 ± 6.38 sec, 5 min; 26.4 ± 6.37 sec | | | | |
| | Latency to 1st interaction (sec.) Target: PPC | ArchT-Light OFF | 10 | 0 | 93.3 ± 24.96 sec | | | | |
| | | ArchT-Light ON | 10 | 0 | 90.0 ± 21.04 sec | One-way ANOVA | F2,24 = 0.008 | P = 0.9920 | N.S. |
| | | EYFP-Light ON | 7 | 0 | 93.7 ± 20.35 sec | | | | |

| | | | | | | | | | |
|---|---|---|---|---|---|---|---|---|---|
| | Interaction time (sec.) Target: BLA | ArchT-Light OFF | 11 | 0 | 1 min; 11.3 ± 5.08 sec, 2 min; 23.7 ± 5.82 sec, 3 min; 29.8 ± 6.12 sec, 4 min; 34.7 ± 4.27 sec, 5 min; 31.2 ± 4.65 sec | Two-way ANOVA with RM | time; $F_{4,128}$ = 1.437<br>group; $F_{4,128}$ = 12.05<br>time x group; $F_{2,32}$ = 1.080 | time; P = 0.1873<br>group; P < 0.0001<br>time x group; P = 0.3515 | time; N.S.<br>group; ****<br>time x group; N.S. |
| | | ArchT-Light ON | 12 | 0 | 1 min; 25.3 ± 3.44 sec, 2 min; 30.3 ± 3.45 sec, 3 min; 35.9 ± 2.83 sec, 4 min; 34.5 ± 2.46 sec, 5 min; 25.3 ± 3.62 sec | | | | |
| | | EYFP-Light ON | 12 | 0 | 1 min; 9.9 ± 4.03 sec, 2 min; 25.5 ± 4.31 sec, 3 min; 29.8 ± 3.92 sec, 4 min; 32.1 ± 3.81 sec, 5 min; 28.8 ± 3.11 sec | | | | |
| | Latency to 1st interaction (sec.) Target: BLA | ArchT-Light OFF | 11 | 0 | 94.4 ± 25.88 sec | One-way ANOVA | $F_{2,32}$ = 4.389 | P = 0.00207 | ** |
| | | ArchT-Light ON | 12 | 0 | 34.1 ± 7.88 sec | | | | |
| | | EYFP-Light ON | 12 | 0 | 97.3 ± 14.60 sec | | | | |
| 3B | Freezing (%) | ChR2-Light OFF in circle | 17 | 0 | Conditioning; 9.5 ± 2.37%, Test; 15.6 ± 2.43% | Two-way ANOVA with RM | session; $F_{1,53}$ = 28.02<br>group; $F_{4,53}$ = 12.53<br>session x group; $F_{4,53}$ = 18.19 | session; P < 0.0001<br>group; P < 0.0001<br>session x group; P < 0.0001 | session; ****<br>group; ****<br>session x group; **** |
| | | ChR2-Light ON in circle | 19 | 0 | Conditioning; 8.6 ± 2.07%, Test; 36.4 ± 4.08% | | | | |
| | | EYFP-Light ON in circle | 9 | 0 | Conditioning; 8.6 ± 2.17%, Test; 11.2 ± 2.23% | | | | |
| | | ChR2-Light OFF in triangle | 7 | 0 | Conditioning; 10.4 ± 4.88%, Test; 7.2 ± 2.70% | | | | |
| | | ChR2-Light ON in triangle | 6 | 0 | Conditioning; 4.5 ± 2.36%, Test; 11.4 ± 3.59% | | | | |
| 4B | Freezing (%) | ArchT-Light OFF | 14 | 0 | Conditioning; 10.1 ± 2.95%, Reactivation; 44.5 ± 3.87%, Test 1; 48.4 ± 2.38%, Test 2; 44.2 ± 2.22% | Two-way ANOVA with RM | session; $F_{3,102}$ = 76.30<br>group; $F_{2,34}$ = 0.4491<br>session x group; $F_{6,102}$ = 4.863 | session; P < 0.0001<br>group; P = 0.6419<br>session x group; P = 0.0002 | session; ****<br>group; N.S.<br>session x group; *** |
| | | ArchT-Light ON | 13 | 0 | Conditioning; 14.7 ± 3.72%, Reactivation; 46.9 ± 3.73%, Test 1; 28.5 ± 4.18%, Test 2; 50.5 ± 2.52% | | | | |
| | | EYFP-Light ON | 10 | 0 | Conditioning; 9.9 ± 2.29%, Reactivation; 40.9 ± 3.89%, Test 1; 43.8 ± 4.14%, Test 2; 42.0 ± 3.83% | | | | |
| 5B | Freezing (%) | ArchT-Light OFF | 11 | 4 | Conditioning; 8.1 ± 1.33%, Reactivation; 45.2 ± 4.40%, Test 1; 50.7 ± 4.04%, Test 2; 49.9 ± 2.95% | Two-way ANOVA with RM | session; $F_{3,96}$ = 87.92<br>group; $F_{2,32}$ = 2.549<br>session x group; $F_{6,96}$ = 4.245 | session; P < 0.0001<br>group; P = 0.0939<br>session x group; P = 0.0008 | session; ****<br>group; N.S.<br>session x group; *** |
| | | ArchT-Light ON | 11 | 4 | Conditioning; 9.6 ± 1.91%, Reactivation; 39.9 ± 2.27%, Test 1; 50.1 ± 4.20%, Test 2; 29.4 ± 4.02% | | | | |
| | | EYFP-Light ON | 13 | 0 | Conditioning; 9.9 ± 1.54%, Reactivation; 43.9 ±4.61%, Test 1; 41.3 ± 4.39%, Test 2; 50.0 ± 2.53% | | | | |
| | Freezing (%) in Test 2 | ArchT-Light OFF | 11 | 4 | 49.9 ± 3.70% | One-way ANOVA | $F_{2,32}$ = 13.64 | P < 0.0001 | **** |
| | | ArchT-Light ON | 11 | 4 | 29.4 ± 5.04% | | | | |
| | | EYFP-Light ON | 13 | 0 | 50.0 ± 3.45% | | | | |
| 5C | Freezing (%) | Vehicle in CA1 | 16 | 0 | Conditioning; 13.9 ± 1.66%, Reactivation; 50.0 ± 2.38%, Test 1; 51.6 ± 2.29%, Test 2; 22.1 ± 2.21% | Two-way ANOVA with RM | session; $F_{3,93}$ = 157.6<br>group; $F_{1,31}$ = 0.9200<br>session x group; $F_{3,93}$ = 0.8081 | session; P < 0.0001<br>group; P = 0.3449<br>session x group; P = 0.4925 | session; ****<br>group; N.S.<br>session x group; N.S. |
| | | Lidocaine in CA1 | 17 | 0 | Conditioning; 15.4 ± 1.92%, Reactivation; 45.3 ± 2.94%, Test 1; 49.1 ± 2.95%, Test 2; 19.2 ± 2.29% | | | | |
| | Motility (pixel / sec.) | Vehicle in CA1 | 16 | 0 | 1-3 min; 82942 ± 5663 pixel, 4-6 min; 52279 ± 2849 pixel, Reactivation; 25391 ± 2195 pixel, Test 1; 20818 ± 1627 pixel, Test 2; 39945 ± 3217 pixel, Test 3; 45339 ± 3250 pixel | Two-way ANOVA with RM | session; $F_{5,155}$ = 87.23<br>group; $F_{1,31}$ = 5.029<br>session x group; $F_{5,155}$ = 6.992 | session; P < 0.0001<br>group; P = 0.0322<br>session x group; P < 0.0001 | session; ****<br>group; *<br>session x group; **** |
| | | Lidocaine in CA1 | 17 | 0 | 1-3 min; 85902 ± 5914 pixel, 4-6 min; 53217 ± 3324 pixel, Reactivation; 25085 ± 1547 pixel, Test 1; 23383 ± 1916 pixel, Test 2; 39270 ± 3184 pixel, Test 3; 79124 ± 6808 pixel | | | | |
| 5D | Freezing (%) | Vehicle in BLA | 13 | 0 | Conditioning; 14.6 ± 2.10%, Reactivation; 49.7 ± 2.94%, Test 1; 50.7 ± 3.29%, Test 2; 22.7 ± 3.04% | Two-way ANOVA with RM | session; $F_{3,72}$ = 105.1<br>group; $F_{1,24}$ = 0.5618<br>session x group; $F_{3,72}$ = 0.8580 | session; P < 0.0001<br>group; P = 0.4608<br>session x group; P = 0.4670 | session; ****<br>group; N.S.<br>session x group; N.S. |
| | | Lidocaine in BLA | 13 | 0 | Conditioning; 13.5 ± 1.72%, Reactivation; 47.9 ± 3.35%, Test 1; 44.5 ± 3.29%, Test 2; 24.0 ± 1.83% | | | | |
| | Interaction time (sec.) | Vehicle in BLA | 13 | 0 | 1 min; 15.4 ± 5.04 sec, 2 min; 24.5 ± 4.71 sec, 3 min; 29.7 ± 4.39 sec, 4 min; 35.4 ± 2.39 sec, 5 min; 31.7 ± 2.64 sec | Two-way ANOVA with RM | time; $F_{4,120}$ = 4.293<br>group; $F_{1,120}$ = 14.01<br>time x group; $F_{4,120}$ = 1.440 | time; P = 0.0028<br>group; P = 0.0003<br>time x group; P = 0.2251 | time; **<br>group;***<br>time x group; N.S. |
| | | Lidocaine in BLA | 13 | 0 | 1 min; 32.5 ± 2.83 sec, 2 min; 35.7 ± 3.17 sec, 3 min; 34.2 ± 2.61 sec, 4 min; 40.6 ± 1.72 sec, 5 min; 34.7 ± 3.66 sec | | | | |
| | Latency to 1st interaction (sec.) | PBS in BLA | 13 | 0 | 77.6 ± 18.87 sec | Unpaired t test | $t_{24}$ = 3.013 | P = 0.0060 | ** |
| | | Lidocaine in BLA | 13 | 0 | 20.1 ± 2.97 sec | | | | |
| 6B | Number of c-Fos positive cells BLA | ArchT-Light OFF | 9 | 0 | 21.3 ± 1.89 cells | One-way ANOVA | $F_{2,29}$ = 11.62 | P = 0.0002 | *** |
| | | ArchT-Light ON | 11 | 0 | 9.9 ± 1.40 cells | | | | |
| | | EYFP-Light ON | 12 | 0 | 19.8 ± 2.02 cells | | | | |
| | Number of c-Fos positive cells ACC | ArchT-Light OFF | 9 | 0 | 23.2 ± 2.49 cells | One-way ANOVA | $F_{2,29}$ = 13.35 | P < 0.0001 | **** |
| | | ArchT-Light ON | 11 | 0 | 8.3 ± 1.05 cells | | | | |
| | | EYFP-Light ON | 12 | 0 | 17.9 ± 2.28 cells | | | | |
| 6D | Number of c-Fos positive cells BLA | ChR2-Light OFF | 11 | 0 | 6.5 ± 0.73 cells | One-way ANOVA | $F_{2,27}$ = 25.44 | P < 0.0001 | **** |
| | | ChR2-Light ON | 12 | 0 | 14.9 ± 1.44 cells | | | | |
| | | EYFP-Light ON | 7 | 0 | 4.0 ± 0.70 cells | | | | |
| | Number of c-Fos positive cells ACC | ChR2-Light OFF | 11 | 0 | 5.6 ± 0.66 cells | One-way ANOVA | $F_{2,27}$ = 12.17 | P = 0.0002 | *** |
| | | ChR2-Light ON | 12 | 0 | 15.8 ± 2.28 cells | | | | |
| | | EYFP-Light ON | 7 | 0 | 5.9 ± 1.32 cells | | | | |
| 6F | Number of c-Fos positive cells BLA | ArchT-Light OFF | 11 | 4 | 22.5 ± 1.70 cells | One-way ANOVA | $F_{2,32}$ = 11.88 | P = 0.0001 | *** |
| | | ArchT-Light ON | 11 | 4 | 12.3 ± 1.24 cells | | | | |
| | | EYFP-Light ON | 13 | 0 | 19.6 ± 1.47 cells | | | | |
| | Number of c-Fos positive cells ACC | ArchT-Light OFF | 11 | 4 | 16.8 ± 2.87 cells | One-way ANOVA | $F_{2,32}$ = 5.096 | P = 0.0120 | * |
| | | ArchT-Light ON | 11 | 4 | 7.6 ± 2.23 cells | | | | |
| | | EYFP-Light ON | 13 | 0 | 16.4 ± 1.71 cells | | | | |
| 7C | Freezing (%) | Light OFF | 12 | 1 | Conditioning; 10.0 ± 1.35%, Test; 54.2 ± 3.20% | Two-way ANOVA with RM | session; $F_{1,23}$ = 140.8<br>group; $F_{1,23}$ = 7.873<br>session x group; $F_{2,23}$ = 18.16 | session; P < 0.0001<br>group; P = 0.0100<br>session x group; P = 0.0003 | session; ****<br>group; *<br>session x group; *** |
| | | Light ON | 13 | 1 | Conditioning; 12.2 ± 1.41%, Test; 33.1 ± 4.73% | | | | |
| 7D | Number of c-Fos positive cells in BLA | Light OFF | 12 | 1 | 29.7 ± 2.16 cell | Unpaired t test | $t_{23}$ = 9.414 | P < 0.0001 | **** |
| | | Light ON | 13 | 1 | 11.5 ± 1.15 cell | | | | |
| | Number of c-Fos positive cells in ACC | Light OFF | 12 | 1 | 44.6 ± 4.13 cell | Unpaired t test | $t_{23}$ = 8.561 | P < 0.0001 | **** |
| | | Light ON | 13 | 1 | 13.4 ± 1.13 cell | | | | |
