## Supplementary Table2 for "Cortical cell ensemble control of past experience-dependent memory updating"

**Table S2. Sampling and statistical analysis, Related to Figure S1-S5 and S7.**

| Fig. # | Category | Group | Sample size (n) Exact size (n) | Sample size (n) Excluded (n) | Score | Statistical test | Degree of Freedom & F/t Value | p-value | Significance? |
|---|---|---|---|---|---|---|---|---|---|
| 1 | Number of Arc+ cells in BLA | Unpaired | 3 | 0 | Cyto; 11.9 ± 0.62, Nuc; 8.4 ± 0.91, Cyto/Nuc; 3.7 ± 0.51 cells | Unpaired t test | BLA;<br>Cyto; t4 = 1.799<br>Nuc; t4 = 0.9879<br>Cyto/Nuc; t4 = 5.376<br><br>CA1<br>Cyto; t4 = 1.009<br>Nuc; t4 = 2.031<br>Cyto/Nuc; t4 = 12.37<br><br>Entorhinal cortex<br>Cyto; t4 = 3.044<br>Nuc; t4 = 2.563<br>Cyto/Nuc; t4 = 5.343<br><br>PFC<br>Cyto; t4 = 3.625<br>Nuc; t4 = 2.415<br>Cyto/Nuc; t4 = 4.500<br><br>Piriform cortex<br>Cyto; t4 = 0.9829<br>Nuc; t4 = 1.677<br>Cyto/Nuc; t4 = 2.734<br><br>Ectorhinal cortex<br>Cyto; t4 = 0.4335<br>Nuc; t4 = 0.5007<br>Cyto/Nuc; t4 = 0.5695<br><br>ACC<br>Cyto; t4 = 6.614<br>Nuc; t4 = 2.261<br>Cyto/Nuc; t4 = 6.002<br><br>Sensory cortex<br>Cyto; t4 = 1.537<br>Nuc; t4 = 1.035<br>Cyto/Nuc; t4 = 1.578<br><br>Visual cortex<br>Cyto; t4 = 1.465<br>Nuc; t4 = 1.094<br>Cyto/Nuc; t4 = 2.150<br><br>Retrosplenial cortex<br>Cyto; t4 = 0.5742<br>Nuc; t4 = 1.594<br>Cyto/Nuc; t4 = 1.112 | BLA;<br>Cyto; P = 0.1464<br>Nuc; P = 0.3791<br>Cyto/Nuc; P = 0.0058<br><br>CA1<br>Cyto; P = 0.3701<br>Nuc; P = 0.1121<br>Cyto/Nuc; P = 0.0002<br><br>Entorhinal cortex<br>Cyto; P = 0.0383<br>Nuc; P = 0.0624<br>Cyto/Nuc; P = 0.0059<br><br>PFC<br>Cyto; P = 0.0223<br>Nuc; P = 0.0731<br>Cyto/Nuc; P = 0.0108<br><br>Piriform cortex<br>Cyto; P = 0.3813<br>Nuc; P = 0.168<br>Cyto/Nuc; P = 0.0522<br><br>Ectorhinal cortex<br>Cyto; P = 0.6870<br>Nuc; P = 0.6429<br>Cyto/Nuc; P = 0.5995<br><br>ACC<br>Cyto; P = 0.0027<br>Nuc; P = 0.0866<br>Cyto/Nuc; P = 0.0039<br><br>Sensory cortex<br>Cyto; P = 0.1992<br>Nuc; P = 0.3591<br>Cyto/Nuc; P = 0.1897<br><br>Visual cortex<br>Cyto; P = 0.2168<br>Nuc; P = 0.3354<br>Cyto/Nuc; P = 0.0979<br><br>Retrosplenial cortex<br>Cyto; P = 0.5966<br>Nuc; P = 0.1861<br>Cyto/Nuc; P = 0.3285 | BLA;<br>Cyto; N.S.<br>Nuc; N.S.<br>Cyto/Nuc; **<br><br>CA1<br>Cyto; N.S.<br>Nuc; N.S.<br>Cyto/Nuc; ***<br><br>Entorhinal cortex<br>Cyto; *<br>Nuc; N.S.<br>Cyto/Nuc; **<br><br>PFC<br>Cyto; *<br>Nuc; N.S.<br>Cyto/Nuc; *<br><br>Piriform cortex<br>Cyto; N.S.<br>Nuc; N.S.<br>Cyto/Nuc; N.S.<br><br>Ectorhinal cortex<br>Cyto; N.S.<br>Nuc; N.S.<br>Cyto/Nuc; N.S.<br><br>ACC<br>Cyto; **<br>Nuc; N.S.<br>Cyto/Nuc; **<br><br>Sensory cortex<br>Cyto; N.S.<br>Nuc; N.S.<br>Cyto/Nuc; N.S.<br><br>Visual cortex<br>Cyto; N.S.<br>Nuc; N.S.<br>Cyto/Nuc; N.S.<br><br>Retrosplenial cortex<br>Cyto; N.S.<br>Nuc; N.S.<br>Cyto/Nuc; N.S. |
| | | Paired | 3 | 0 | Cyto; 15.2 ± 1.75, Nuc; 9.4 ± 0.44, Cyto/Nuc; 7.4 ± 0.48 cells | | | | |
| | Number of Arc+ cells in CA1 | Unpaired | 3 | 0 | Cyto; 28.7 ± 2.85, Nuc; 13.0 ± 1.07, Cyto/Nuc; 7.3 ± 0.19 cells | | | | |
| | | Paired | 3 | 0 | Cyto; 32.2 ± 2.08, Nuc; 16.7 ± 1.45, Cyto/Nuc; 13.8 ± 0.48 cells | | | | |
| | Number of Arc+ cells in Entorhinal cortex | Unpaired | 3 | 0 | Cyto; 22.6 ± 0.40, Nuc; 11.4 ± 1.09, Cyto/Nuc; 6.4 ± 0.48 cells | | | | |
| | | Paired | 3 | 0 | Cyto; 27.4 ± 1.56, Nuc; 15.2 ± 0.99, Cyto/Nuc; 13.7 ± 1.26 cells | | | | |
| | Number of Arc+ cells in Piriform cortex | Unpaired | 3 | 0 | Cyto; 11.9 ± 1.16, Nuc; 6.7 ± 0.67, Cyto/Nuc; 3.9 ± 0.73 cells | | | | |
| | | Paired | 3 | 0 | Cyto; 13.7 ± 1.39, Nuc; 10.3 ± 2.08, Cyto/Nuc; 6.2 ± 0.44 cells | | | | |
| | Number of Arc+ cells in Prefrontal cortex | Unpaired | 3 | 0 | Cyto; 20.2 ± 1.46, Nuc; 8.4 ± 1.11, Cyto/Nuc; 5.7 ± 1.15 cells | | | | |
| | | Paired | 3 | 0 | Cyto; 26.7 ± 1.02, Nuc; 13.2 ± 1.64, Cyto/Nuc; 11.7 ± 0.67 cells | | | | |
| | Number of Arc+ cells in Anterior cingurate cortex | Unpaired | 3 | 0 | Cyto; 11.3 ± 0.19, Nuc; 5.8 ± 0.29, Cyto/Nuc; 3.1 ± 0.56 cells | | | | |
| | | Paired | 3 | 0 | Cyto; 15.2 ± 0.56, Nuc; 7.4 ± 0.68, Cyto/Nuc; 7.0 ± 0.33 cells | | | | |
| | Number of Arc+ cells in Ectorhinal cortex | Unpaired | 3 | 0 | Cyto; 26.2 ± 3.25, Nuc; 11.0 ± 1.26, Cyto/Nuc; 7.7 ± 1.07 cells | | | | |
| | | Paired | 3 | 0 | Cyto; 28.2 ± 3.28, Nuc; 12.8 ± 3.32, Cyto/Nuc; 9.0 ± 2.08 cells | | | | |
| | Number of Arc+ cells in Visual cortex | Unpaired | 3 | 0 | Cyto; 41.2 ± 5.86, Nuc; 14.3 ± 5.10, Cyto/Nuc; 8.7 ± 2.99 cells | | | | |
| | | Paired | 3 | 0 | Cyto; 50.0 ± 1.26, Nuc; 20.6 ± 2.51, Cyto/Nuc; 16.0 ± 1.64 cells | | | | |
| | Number of Arc+ cells in Sensory cortex | Unpaired | 3 | 0 | Cyto; 37.0 ± 6.08, Nuc; 11.4 ± 1.56, Cyto/Nuc; 9.1 ± 1.16 cells | | | | |
| | | Paired | 3 | 0 | Cyto; 46.9 ± 2.12, Nuc; 15.2 ± 3.30, Cyto/Nuc; 13.3 ± 2.41 cells | | | | |
| | Number of Arc+ cells in Retrosplenial cortex | Unpaired | 3 | 0 | Cyto; 22.3 ± 3.67, Nuc; 9.0 ± 1.67, Cyto/Nuc; 7.0 ± 1.50 cells | | | | |
| | | Paired | 3 | 0 | Cyto; 25.7 ± 4.50, Nuc; 5.7 ± 1.26, Cyto/Nuc; 4.9 ± 1.16 cells | | | | |
| 2C | Interaction time (sec.) | No-IS | 12 | 0 | 1 min; 36.4 ± 3.82 sec, 2 min; 31.6 ± 4.19 sec, 3 min; 32.7 ± 3.71 sec,<br>4 min; 31.6 ± 4.50 sec, 5 min; 24.5 ± 4.83 sec | Two-way ANOVA with RM | time; F4,110 = 0.9164<br>group; F1,110 = 15.93<br>time x group; F4,110 = 3.369 | time; P = 4572<br>group; P = 0.0001<br>time x group; P = 0.0122 | time; N.S.<br>group; ***<br>time x group; * |
| | | IS | 12 | 0 | 1 min; 12.2 ± 4.19 sec, 2 min; 21.1 ± 4.11 sec, 3 min; 27.0 ± 3.73 sec,<br>4 min; 26.1 ± 4.99 sec, 5 min; 25.1 ± 5.56 sec | | | | |
| | Latency to 1st interaction (sec.) | No-IS | 12 | 0 | 7.4 ± 1.22 sec | Unpaired t test | t22 = 4.234 | P = 0.0003 | *** |
| | | IS | 12 | 0 | 446.9 ± 9.25 sec | | | | |
| 3 | Zif268 positive cells / DAPI (%) | OFF Dox ArchT-Light OFF | 6 | 0 | 38.2 ± 2.76% | One-way ANOVA | F3,20 = 15.02 | P < 0.0001 | **** |
| | | OFF Dox ArchT-Light ON | 6 | 0 | 14.1 ± 2.22% | | | | |
| | | ON Dox ArchT-Light ON | 6 | 0 | 35.7 ± 4.00% | | | | |
| | | OFF Dox EYFP-Light ON | 6 | 0 | 42.6 ± 3.78% | | | | |
| 4A | Freezing (%) | Vehicle | 11 | 6 | Conditioning; 8.3 ± 3.16%, Test; 45.1 ± 5.53% | Two-way ANOVA with RM | session; F1,22 = 57.14<br>group; F1,22 = 13.02<br>session x group; F1,22 = 17.83 | session; P < 0.0001<br>group; P = 0.0016<br>session x group; P = 0.0004 | session; ****<br>group; **<br>session x group; *** |
| | | Lidocaine | 13 | 7 | Conditioning; 7.0 ± 2.52%, Test; 17.4 ± 2.92% | | | | |
| 4B | Motility (pixel/min) | Vehicle | 7 | 1 | Conditioning;<br>1 min; 80233 ± 6772 pixel, 2 min; 70457 ± 6243 pixel, 3 min; 71208 ± 7504 pixel,<br>4 min; 65571 ± 7484 pixel, 5 min; 60323 ± 5282 pixel, 6 min; 52383 ± 6142 pixel<br><br>Test;<br>1 min; 62408 ± 8760 pixel, 2 min; 47422 ± 7876 pixel, 3 min; 55392 ± 6786 pixel | Two-way ANOVA with RM | Conditioning session;<br>time; F5,78 = 8.644<br>group; F1,78 = 0.2657<br>time x group; F5,78 = 0.7214<br><br>Test session;<br>time; F2,39 = 2.946<br>group; F1,39 = 1.382<br>time x group; F2,39 = 0.5359 | Conditioning session;<br>time; P < 0.0001<br>group; P = 0.6077<br>time x group; P = 0.6094<br><br>Test session;<br>time; P = 0.0643<br>group; P = 0.2469<br>time x group; P = 0.5894 | Conditioning session;<br>time; ****<br>group; N.S.<br>time x group; N.S.<br><br>Test session;<br>time; N.S.<br>group; N.S.<br>time x group; N.S. |
| | | Lidocaine | 8 | 1 | Conditioning;<br>1 min; 89938 ± 6206 pixel, 2 min; 80387 ± 5205 pixel, 3 min; 75371 ± 4348 pixel,<br>4 min; 59187 ± 6464 pixel, 5 min; 55224 ± 5323 pixel, 6 min; 50793 ± 4843 pixel<br><br>Test;<br>1 min; 59146 ± 4422 pixel, 2 min; 46064 ± 6044 pixel, 3 min; 41967 ± 3082 pixel | | | | |
| | | Vehicle-No IS | 8 | 1 | 1 min; 23.4 ± 4.53 sec, 2 min; 25.7 ± 4.69 sec, 3 min; 31.1 ± 3.36 sec,<br>4 min; 35.2 ± 2.85 sec, 5 min; 30.6 ± 3.72 sec | | | | |

| | | | | | | | | | |
|---|---|---|---|---|---|---|---|---|---|
| 4C | Interaction time (sec.) | Vehicle-IS | 8 | 2 | 1 min; 7.8 ± 4.00 sec, 2 min; 18.8 ± 5.49 sec, 3 min; 30.3 ± 3.06 sec, 4 min; 26.7 ± 3.74 sec, 5 min; 29.5 ± 4.13 sec | Two-way ANOVA with RM | time; F4,88 = 14.80<br>group; F2,22 = 1.508<br>time x group; F8,88 = 1.347 | time; P < 0.0001<br>group; P = 0.2434<br>time x group; P = 0.2314 | time; ****<br>group; N.S.<br>time x group; N.S. |
| | | Lidocaine-IS | 9 | 2 | 1 min; 8.6 ± 3.29 sec, 2 min; 23.7 ± 4.57 sec, 3 min; 25.4 ± 3.80 sec, 4 min; 31.3 ± 5.29 sec, 5 min; 33.2 ± 3.63 sec | | | | |
| | Latency to 1st interaction (sec.) | Vehicle-No IS | 8 | 1 | 27.5 ± 10.29 sec | One-way ANOVA | F2,22 = 4.626 | P = 0.0210 | * |
| | | Vehicle-IS | 8 | 2 | 80.4 ± 16.36 sec | | | | |
| | | Lidocaine-IS | 9 | 2 | 78.3 ± 13.83 sec | | | | |
| 5 | c-Fos positive cells / DAPI (%) | OFF Dox ChR2-Light OFF | 6 | 0 | 0.3 ± 0.19% | One-way ANOVA | F3,20 = 779.7 | P < 0.0001 | **** |
| | | OFF Dox ChR2-Light ON | 6 | 0 | 20.0 ± 0.58% | | | | |
| | | ON Dox ChR2-Light ON | 6 | 0 | 0.1 ± 0.09% | | | | |
| | | OFF Dox EYFP-Light ON | 6 | 0 | 0.6 ± 0.34% | | | | |
| 7 | Zif268 positive cells / DAPI (%) | OFF Dox Light OFF | 6 | 0 | 13.5 ± 1.25% | One-way ANOVA | F2,15 = 54.14 | P < 0.0001 | **** |
| | | OFF Dox Light ON | 6 | 0 | 2.0 ± 0.21% | | | | |
| | | ON Dox Light ON | 6 | 0 | 10.7 ± 0.63% | | | | |
